# Generation of surrogate brain maps preserving spatial autocorrelation through random rotation of geometric eigenmodes

**DOI:** 10.1101/2024.02.07.579070

**Authors:** Nikitas C. Koussis, James C. Pang, Jayson Jeganathan, Bryan Paton, Alex Fornito, P. A. Robinson, Bratislav Misic, Michael Breakspear

## Abstract

The brain expresses activity in complex spatiotemporal patterns, reflected in the influence of spatially distributed cytoarchitectural, biochemical, and genetic properties. The correspondence between these multimodal “brain maps” may reflect underlying causal pathways and is hence a topic of substantial interest. However, these maps possess intrinsic smoothness (spatial autocorrelation, SA) which can inflate spurious cross-correlations, leading to false positive associations. Identifying true associations requires knowledge about the distribution of correlations that arise by chance in the presence of SA. This null distribution can be generated from an ensemble of surrogate brain maps that preserve the intrinsic SA but break the correlations between maps. The present work introduces the “eigenstrapping” method, which uses a spectral decomposition of cortical and subcortical surfaces in terms of geometric eigenmodes, and then randomly rotating these modes to produce SA-preserving surrogate brain maps. It is shown that these surrogates appropriately represent the null distribution of chance pairwise correlations, with similar or superior false positive control to current state-of-the-art procedures. Eigenstrapping is fast, eschews the need for parametric assumptions about the nature of a map’s SA, and works with maps defined on smooth surfaces with or without a boundary. Moreover, it generalizes to broader classes of null models than existing techniques, offering a unified approach for inference on cortical and subcortical maps, spatiotemporal processes, and complex patterns possessing higher-order correlations.

## MAIN TEXT

Interest in spatial patterns of cortical activity, cellular and microstructural composition, molecular architecture, and network connectivity of the brain has surged in recent years ^1–6^. An important challenge in this field is to measure the similarity between two or more such “brain maps” while excluding spurious relationships arising from chance. Correlations between different maps may reflect the influence of spatially patterned gene expression on cytoarchitecture or neuronal activity, hence motivating further mechanistic investigation ^3,7^. However, cortical regions that are close together tend to possess similar features, the causes of which may be biological (such as a gradual change in gene expression) or methodological (due to the spatial smoothing that is applied in the analyses of most imaging modalities). These effects combine to endow brain maps with a spatial autocorrelation (SA) that typically has an extent of tens of mm ^5,8–10^. The presence of such within-map correlations reduces the true degrees of freedom when testing for pairwise associations between maps, hence amplifying spurious associations. Null hypotheses of the correspondence between maps, i.e., the distribution of “chance” in map-to-map correlations, need to preserve SA to control Type I error ^8,11^. This is not a trivial undertaking in the presence of complex statistical dependencies within and between maps ^12^.

There are several methods that can generate surrogate maps that maintain SA while randomizing the association between maps, hence providing suitable “null models”. Most of these null models fall into two broad classes: 1) direct spatial permutation, commonly known as the “Spin Test” ^10,13–16^, whereby maps in the neocortex are projected onto a sphere, randomly rotated, then projected back to the cortical surface; and 2) parameterized spatial randomization, such as “Brain Surrogate Maps with Autocorrelated Spatial Heterogeneity” (BrainSMASH), whereby surrogate maps are drawn from a random (Gaussian) process and smoothed to match the empirical SA with parametric models that approximate the original statistical structure ^8,17^. However, both classes have drawbacks: the Spin Test provides incomplete coverage of the cortex because it rotates missing data in the medial wall (i.e., vertices within the subcortex and anatomically inferior to the cingulate) onto the map (see Fig. 5 for a demonstration). In addition, the Spin Test has thus far not been extended to volumetric maps, precluding its use in the subcortex. The Spin Test also preserves the original spatial relationships between all points, only rotating them to different locations. This form of randomization yields a restricted null space with an assumption that no higher-order spatial structure exists within the original map. Higher-order spatial effects occur frequently in biological systems including neural processes in visual cortex, which reflect the complex statistical dependences in natural scenes ^18–21^. Estimating the null space to identify these more complex spatial effects requires a randomization of higher-order statistical dependencies. Conversely, generating spatial nulls with spatial parametric techniques such as BrainSMASH ^8^ requires extensive parameter optimization and is computationally intensive ^11^. Moreover, these methods rest upon assumptions of stationarity on brain maps, drawing randomness from stationary Gaussian processes. Cortical activity frequently violates these assumptions, exhibiting long-tailed statistics and nonlinear spatiotemporal properties ^22–27^. At high levels of SA, both of these methods fail tests of Type I error, with false positive rates 2–10 times higher than expected ^11^. This inflation can be particularly problematic for inference on very smooth, lower-resolution maps, such as those generated with brain transcriptomics or positron emission tomography.

To improve on the current state of null models, we turn to geometric basis sets. These basis sets – known as geometric eigenmodes – support the decomposition of complex spatial patterns from coarse to fine wavelengths. Geometry constrains the behavior of many complex systems, including the brain, where it influences large-scale dynamics such as standing and travelling waves ^28–32^. Geometric eigenmodes have increasingly been used to model and describe these diverse aspects of brain activity and structure ^33–39^. Geometric basis sets are essentially spherical harmonics generalized to non-spherical surfaces and can be derived by application of the Laplace-Beltrami operator (LBO) ^34^. Notably, for the present purposes, the LBO projects spatial data into an orthogonal subspace where the data representation (the eigenmode coefficients) are decorrelated and hence exchangeable, similar to the Fourier or wavelet transforms ^40,41^. This allows constrained randomization of spatial data on irregular surfaces without disrupting (two-point) spatial correlations when the data are back-projected into the original spatial domain. Appropriate eigenmode randomization can thus yield a geometric surrogate map preserving the SA of the original data while randomizing the location and higher-order properties of the map.

Here, we introduce *eigenstrapping*, a method of generating random brain maps with preserved SA for null hypothesis testing. By leveraging the mathematical properties of the LBO, eigenstrapping provides a method to perform rigorous statistical inference of cortical and subcortical associations and surface or volumetric maps for a broad range of research questions. We show that eigenstrapping has distinct advantages over existing methods for producing surrogate brain maps, including stronger false positive rate (FPR) control, relatively low computational burden, generalizability to a broad class of spatial processes, use in both cortical and subcortical maps, and applicability to complex spatial and spatiotemporal processes ^10^. We provide an open-source Python-based package that is deployable to commonly utilized neuroimaging formats ^42^.

## RESULTS

We first describe the method for decomposing a brain map into cortical eigenmodes. The method of constrained randomization through rotation of these eigenmodes is then provided. We next show how reconstructing a map from these rotated modes produces SA-preserving surrogate maps. Ensembles of these surrogates obtained through repeated random eigenmode rotation yields a null distribution for spurious associations between smooth maps, which we benchmark against the Spin Test and BrainSMASH methods. We end with an exposition of the randomization of more complex (ternary and quaternary) correlations by eigenmode rotation and the relevance of this to probe brain maps for complex textural properties.

### Eigenmode decomposition and group-based rotation

An eigenmode decomposition on a discretized surface *x* with *N* vertices yields *N*-1 orthogonal eigenmodes which can be ordered by their corresponding eigenvalues. These modes allow a spectral decomposition of a spatial pattern *y* from coarse to fine wavelengths (i.e., spatial frequencies) ^33,34,43^. In the spherical case, the eigenmodes are called spherical harmonics and occur in groups of modes with identical (degenerate) eigenvalues. Each harmonic group thus describes a set of orthogonal spatial patterns expressed on the sphere that have the same characteristic wavelength. The folds, gyri, and non-spherical distortions of the cortical geometry perturb this structure, but the eigenvalue separation between groups of modes is approximately preserved, particularly at spatial scales relevant for whole brain maps, allowing one to use similar groupings ^33,34^.

Formally, an empirical brain map *y*(*x*) on a discrete surface *x* with *N* vertices is decomposed into a linear combination of geometric eigenmodes,

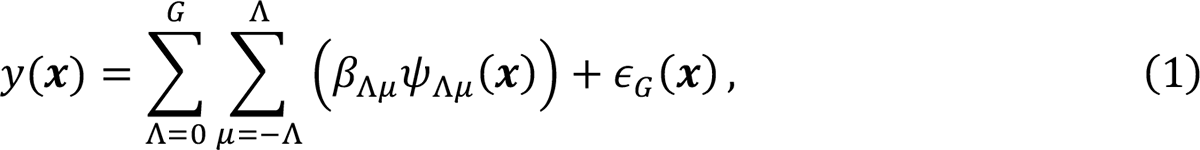

where *G* is the total number of groups used in the decomposition and there are 2Λ + 1 modes in each group. β_Λμ_ is the linear coefficient (weighting) of mode ψ_Λμ_ in group Λ with eigenvalue λ_Λμ_ (see Fig. 1A). These coefficients are estimated by integration of the modes with the data *y* on the surface ***x*** (see Methods) ^44,45^. The residual error ∈_*G*_(*x*) decreases in amplitude as the number of groups *G* used in the decomposition increases, vanishing if the decomposition is complete; that is, if the full complement of *N* − 1 modes is used. Individual modes within a group Λ are orthogonal by virtue of their relative orientation, whilst the groups themselves are also orthogonal due to their differing characteristic wavelengths. We use this orthogonality between groups to resample modes without disrupting the spatial spectra and hence smoothness of the map *y*. A more detailed description is provided in Methods.

**Fig. 1.**
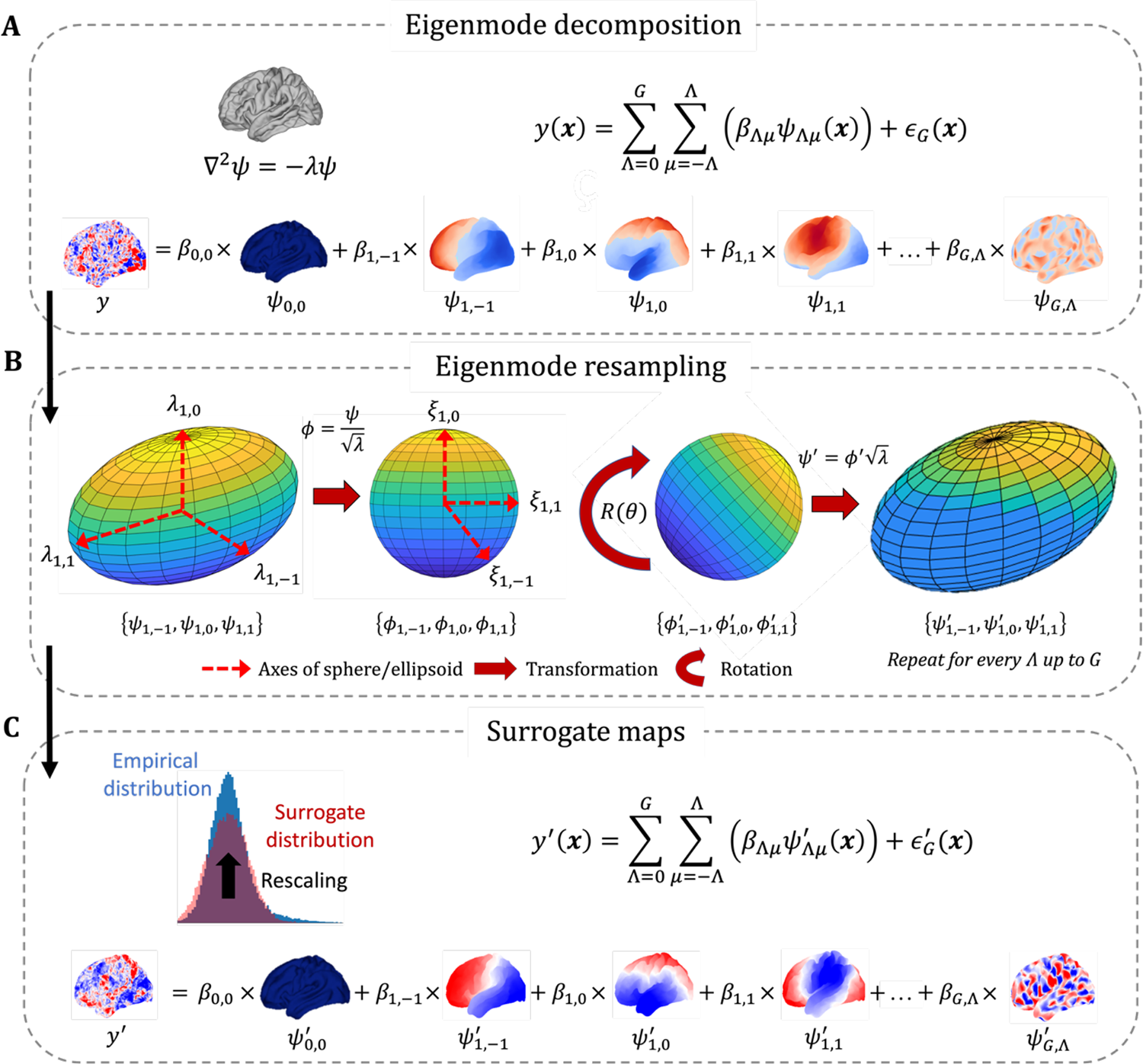
The *eigenstrapping* method to generate surrogates that preserve spatial autocorrelation. (A) *Eigenmode decomposition*: coefficients β_Λμ_ are derived from the generalized linear model (GLM; Eq. 1). A total number of modes is chosen such that the residual error in the GLM is negligible. (B) *Eigenmode rotation*: Eigenmodes are partitioned into eigengroups Λ (of size *n* = 2Λ + 1) and normalized by their eigenvalues λ_Λμ_ to yield spherical eigenmodes Ø_Λμ_ with identical eigenvalues ξ_Λμ_. This is analogous to transformation from an *n*-dimensional ellipsoid with axes λ_Λμ_to an *n*-dimensional sphere with axes ξ_Λμ_. The equality of ξ (i.e., degeneracy; Fig. S1) permits rotation of Ø_Λμ_ by a random rotation matrix *R*(θ), resulting in rotated spherical eigenmodes Ø^’^. These modes are multiplied by /λ_Λμ_ to project them back to the ellipsoid, resulting in rotated modes ψ^’^ in groups Λ. (C) *Surrogate maps*: The GLM with original coefficients β is multiplied by rotated modes ψ^’^ across all Λμ yielding a surrogate brainmap *y*^’^. An optional amplitude adjustment step (*rescaling*; see Supplementary Information-S2) is applied to the reconstructed data (in red; original data in blue). Residuals can be permuted and added back into the resulting surrogate map *y*^’^ (see Supplementary Information-S1).

Spherical harmonics within groups possess identical characteristic spatial frequencies (eigenvalues) while the groups themselves are invariant under rotation. Geometric eigenmodes adapt to the folds and undulations of the cortical surface. As a consequence, geometric mode groups are not rotationally invariant and modes within a group possess similar but not identical spatial frequencies. To rotate geometric eigenmodes within groups, it is thus necessary to normalize their eigenvalues to have equal value, equivalent to mapping the modes onto an *n*-dimensional sphere *Z*,

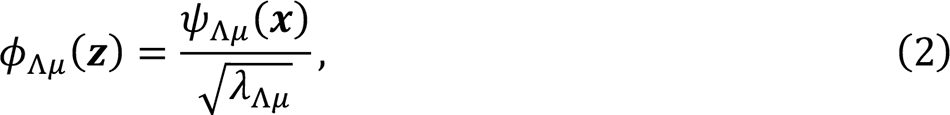

where Ø_Λμ_ is the equivalent spherical representation of ψ_Λμ_. The number of modes in a group and hence the dimension of the sphere remains *n* = 2Λ + 1 (Supplementary Table S1).

As a result of this normalization, all spherical modes within a group have identical eigenvalues denoted ξ_Λ1_ = ξ_Λ2_ = ⋯ ξ_Λ*n*_. To perform eigenstrapping, groups of spherical modes are rotated by taking the matrix multiplication with a random rotation matrix *R*(θ_Λ_) ^46^,

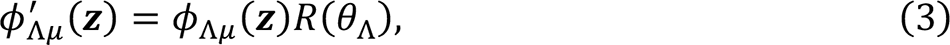

where the prime denotes rotation by a random angle θ and Λ is the group number. This group-based rotation ensures that spherical modes within groups retain their orthogonality. This process is repeated with an independent random rotation applied to each group, breaking the original angular alignment of modes in different groups.

Rotated spherical modes Ø* (*Z*) are then mapped back to the cortical geometry, yielding rotated geometric eigenmodes,

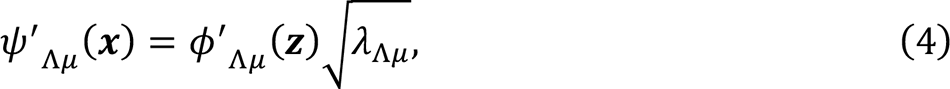

A surrogate brain map *y*\*(*x*) is then obtained from these rotated eigenmodes,

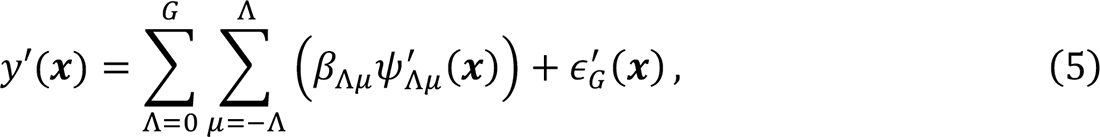

where β_Λμ_ are the same coefficients from Eq. (1) (Fig. 1C). The surrogate error term ∈* (*x*) is derived from simple random permutation of the error term ∈_*G*_(*x*) from Eq. (1) (see Methods and Supplementary Information-S1). To preserve the amplitude distribution of the empirical data, an optional amplitude-adjustment step can be performed (Fig. 1C – top left; Supplementary Information-S2). The resampling procedure is illustrated in Fig. 1B for the first non-zero eigengroup, Λ = 1.

### Statistical properties of surrogate maps generated from rotated eigenmodes

To test the performance of this procedure, we generated eigenstrapping surrogate maps of task-evoked fMRI data on the *fs-LR-32k* surface of 255 unrelated healthy individuals from the Human Connectome Project ^47^ (HCP; *emotion*, see Supplementary Table S2 for a list of tasks). This was compared to surrogates generated from SA-naïve random permutation of vertices. An example target map (HCP; *gambling*) is compared to an example contrast *emotion* map in Fig. 2A with Pearson’s correlation *r* = 0.249. Example surrogate maps using an eigenmode decomposition with 6000 modes visually capture the smoothness of the original data (Fig. 2B). Quantifying the SA of these maps using the variogram, a measure of local smoothness ^8^ shows that eigenstrapping (Fig. 2C, blue) preserves the empirical SA to very small spatial separations (<1 mm). In contrast, SA-naïve random permutations whiten the surrogates, producing relatively flat SA (Fig. 2C, red). An ensemble of 1000 eigenstrapped surrogate maps exhibits a broad, zero-centered distribution of correlations with a target empirical map (Fig. 2D, blue; *gambling*), hence yielding a wider distribution than SA-naïve random permutations (Fig. 2D, red). Notably, eigenstrapped surrogate maps are on average uncorrelated with each other, yielding a broad, zero-centered pairwise-correlation distribution (Fig. 2E).

**Fig. 2.**
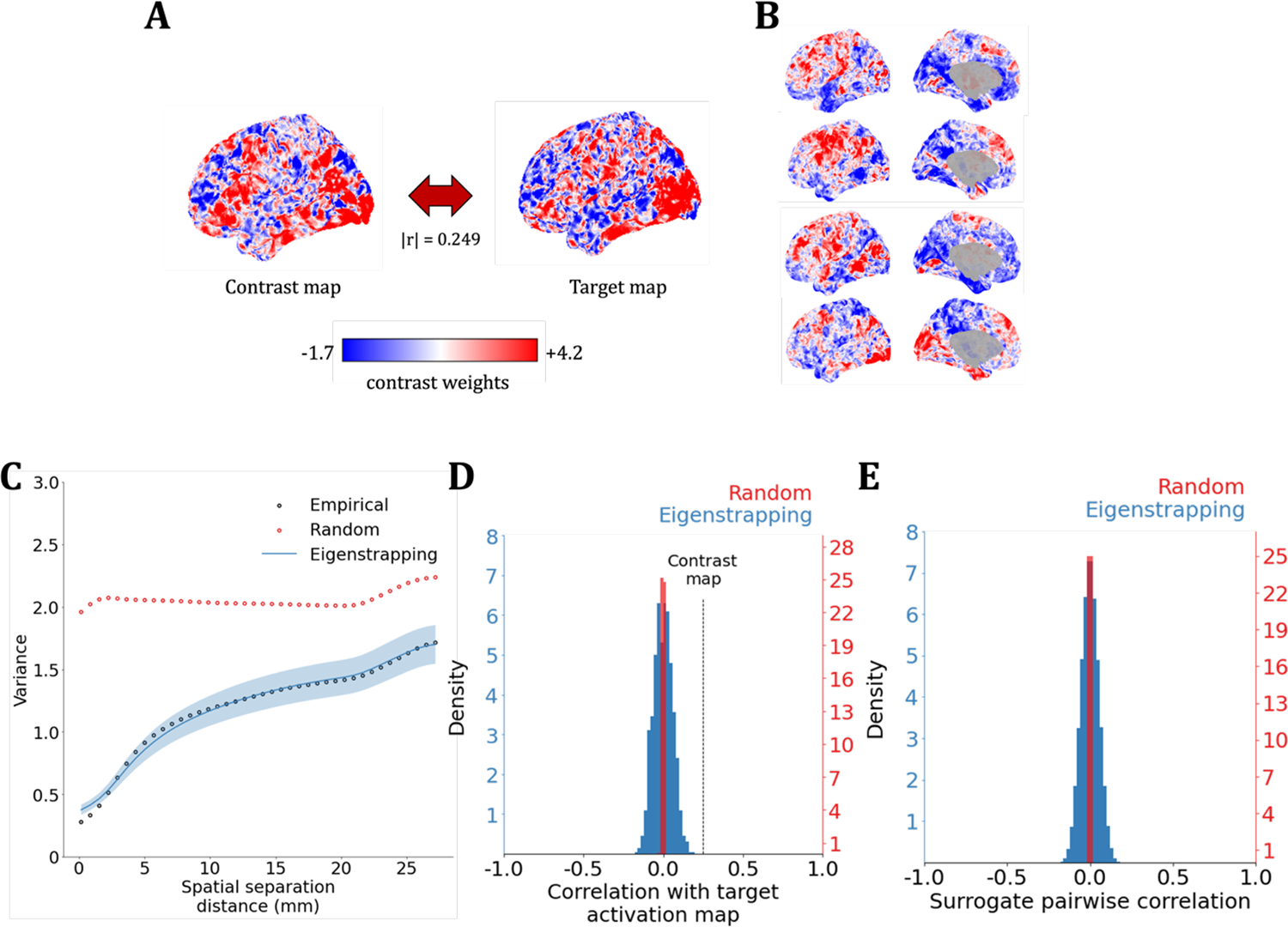
Statistical properties of eigenstrapped surrogates of fMRI data. (A) An example HCP task contrast map (*Contrast map*; left) was correlated with a target contrast map (*Target map*; right) from the same participant in another task condition at |*r*| = 0.249. Each map is colored by task contrast weight. (B) Four example eigenstrapped surrogates of the contrast map (panel A, left). (C) The variogram from 0 to 30 mm spatial separation with the average of 1000 SA-naive surrogates (*Random*; red circles), and the average and standard deviation of 1000 eigenstrapping surrogates (*Eigenstrapping*; blue line and shading, respectively) against the contrast map (*Empirical*). (D) Correlation of SA-naïve (red) and eigenstrapping (blue) surrogates with the target fMRI map. The correlation of the target map to the contrast map |*r*| = 0.249 is shown with the dashed black line. In this case, the correlation lies outside the null distribution and it thus considered statistically significant. (D) Pairwise correlation of SA-naive (red) and eigenstrapping (blue) surrogates.

### Control of false positives in simulated brain maps

We next tested the efficacy of eigenstrapping in controlling Type I error, benchmarked against a ground truth from simulated brain maps. Simulated maps were generated with Gaussian random fields (GRFs) that have parametrically varying SA ^48,49^ with smoothness parameter α (Fig. 3; see Methods and Supplementary Information-S4). We simulated pairs of GRF maps with predetermined cross-correlations of |*r*| = 0.15±0.005 and a cortical resolution of 10,242 vertices in the *fsaverage5* standard space. The relatively weak correlation of |*r*| = 0.15 lies outside the null distribution for pairs of cortical maps that possess weak SA (low α), but falls within the null, consistent with a chance association, for pairs of smooth cortical maps (high α) ^8,11^. Smoothness was tuned from α = 0.0 (no SA) to α = 4.0 (high SA) in steps of 0.5 with 1000 pairs of GRFs generated at each step (Fig. 3A-B; Fig. S4). This procedure yielded 9000 total pairs with pairwise cross-correlations centered at 0.15 (range 0.145-0.155).

**Fig. 3.**
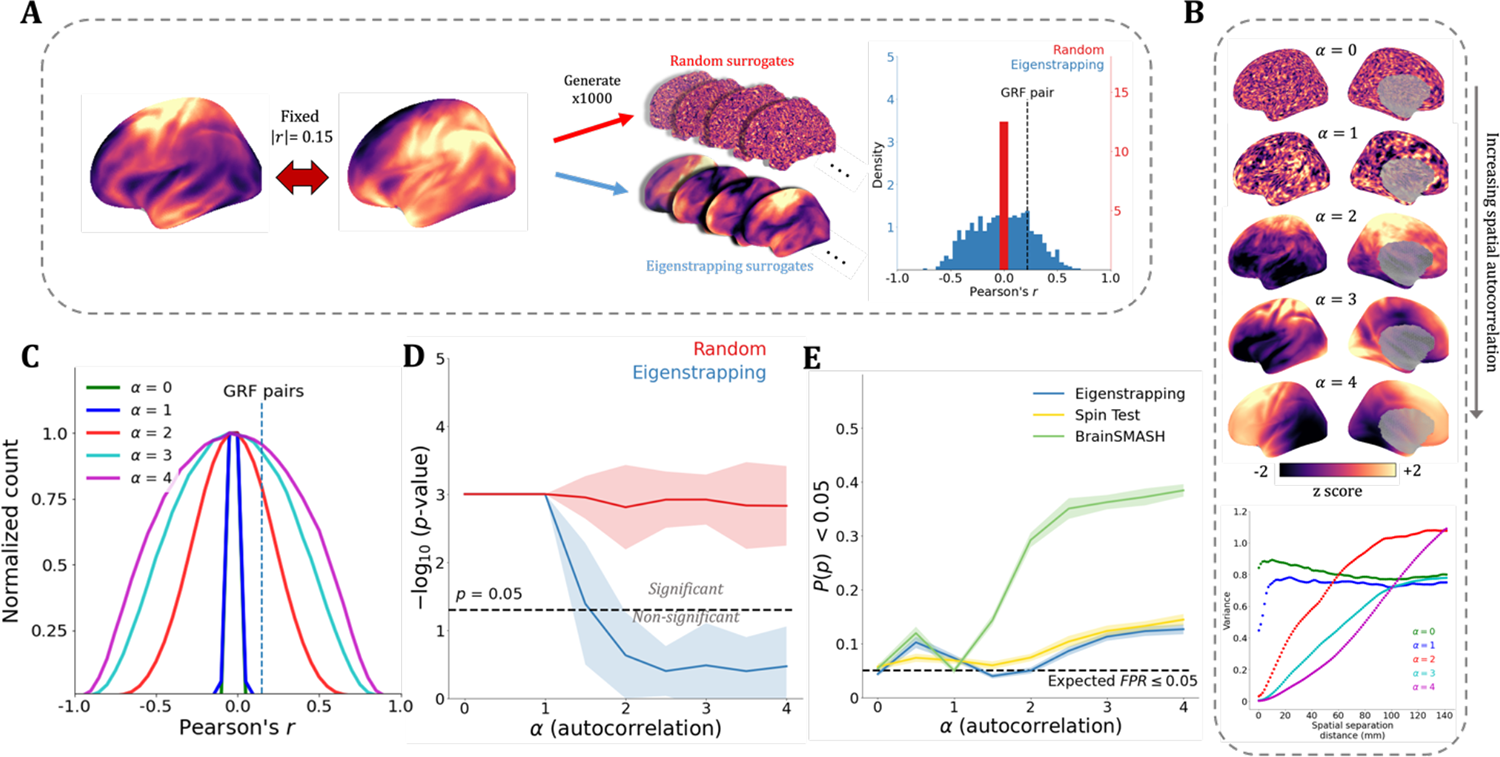
Eigenstrapping control of false positives. (A) Gaussian random fields (GRFs) with varying SA were used to generate pairs of cortical maps with absolute Pearson correlation |*r*| = 0.15±0.005. Two GRF maps are plotted at α = 2 alongside random (red) and eigenstrapping (blue) exemplar surrogates. Rightmost panel shows exemplar histogram of SA-randomizing surrogate (red) and eigenstrapping surrogate (blue) correlations. Dashed line shows the ground-truth correlation of the GRF pair. (B) GRFs are plotted with α increasing from 0.0 to 4.0. Variograms derived from the generated maps demonstrate the increase in SA with increase in α (bottom panel). The choice of modes for eigenstrapping of each α-pair was tailored to the best visual fit of surrogate to empirical variograms. (C) Average null distributions of eigenstrapping for different levels of SA, normalized between 0 and 1. (D) Mean and standard deviation of two-tailed *p*-values of 1000 surrogates (SA-randomizing: red; eigenstrapping: blue) per 9000 GRF pairs as a function of SA. Black dashed line shows significance at *p* = 0.05 (-log_10_(*p*) = 1.3). Null hypotheses are rejected above this line, not rejected below this line. (E) Each line indicates false positive rate (FPR) of null method as a function of spatial autocorrelation (Eigenstrapping: blue; Spin Test: yellow; BrainSMASH: green). Mean FPRs after 20 randomized sets are shown in solid lines. Shaded areas around solid lines correspond to standard deviations. The black dashed line corresponds to expected FPR of ≤ 5%.

We used eigenstrapping to derive a significance value for the cross-correlation of the simulated pairs of cortical maps and compared it to random, SA-naïve surrogates of the same maps. 1000 surrogate maps were derived from one map in each GRF pair using eigenstrapping (Fig. 3A, blue) with fixed numbers of modes based on the SA (Fig. 3B). The eigenstrapped surrogates and the other map in the GRF pair were correlated, forming a correlation distribution across α levels (Fig. 3C). The choice of modes was tailored empirically against the average GRF variogram, which is 2500 at α = 0.0-1.0, 1500 at α = 1.5, 500 at α = 2.0, 200 at α = 2.5-3.5, and 50 at α = 4.0. At high SA (α > 1.5), the variogram is preserved with relatively few modes (corresponding to 0.05–1.5% of all modes on the surface) following amplitude adjustment (Supplementary Information-S2). This was compared to SA-naïve random surrogates, obtained by randomly permuting the data (Figure 2A; red). These distributions were then used to estimate the two-tailed *p*-value for the original correlation of the GRF pair. As SA increases, the distribution of the correlation between eigenstrapped surrogates widens (Fig. 2C) until the inter-map correlation of 0.15 falls within the tail of the distribution at α ≥ 2. The *p*-value increases accordingly (i.e., the -log_10_(*p*) drops) becoming greater than 0.05 for α ≥ 2 (-log_10_(*p*) < 1.3; Fig. 2D). This analysis thus shows that a chance cross-correlation of |*r*| = 0.15 is common among smooth brain maps. The distribution of correlations remains narrow for all randomized surrogates (Fig. 2D; red) and the original correlation remains well outside the distribution for all α (that is, the null is too precisely represented by SA-naïve nulls).

We next assessed the false positive rate (FPR) of eigenstrapping against those of the Spin Test and BrainSMASH (the two most cited methods for spatial null models) by randomly swapping one map from each of the pairs from the analysis in Fig. 2A-D. Since the correlations between these randomly paired maps will be zero centered, the FPR should be equal to or below the chosen statistical alpha – i.e., ≤ 5% FPR at *p* < 0.05. Eigenstrapping yields an FPR close to the expected 5% for low SA (α < 1.5, Fig. 2E). As SA increased to a level visually consistent with the smoothness of empirical brain maps (α ≥ 1.5), eigenstrapping performs below expected at 3.9% (α = 1.5), expected at 5.0% (α = 2.0), then increases to 8.6% (α = 2.5), and 11.3% (α = 3.0). At higher levels of SA (smoother than typically seen empirically), the FPR rises to 12.3% (α = 3.5) and 12.6% (α = 4.0). For the same test, the Spin Test yields slightly higher FPR than eigenstrapping across all high SA regimes (Fig. 2E in yellow) at 5.9% (*a* = 1.5), 7.5% (*a* = 2.0), 10.4% (α = 2.5), 12.3% (α = 3.0), 13.2% (α = 3.5), and 14.5% (α = 4.0). The BrainSMASH method shows much higher FPR than both the Spin Test and eigenstrapping, reaching 29.2% at α = 2.0 and 38.4% at α = 4.0 (Fig. 2E in green).

We further quantify the SA-preserving property of eigenstrapping by calculating Moran’s *I*, a measure of global SA ^50^, for each GRF and surrogate map (Fig. S5-6). In contrast to the variogram, which captures two-point correlations as a function of distance, Moran’s *I* provides a single composite summary of SA ^8,50^. Eigenstrapping preserves Moran’s *I* for all levels of smoothness α (see Supplementary Information-S5).

### Null hypothesis testing of associations between empirical brain maps

A primary goal of using surrogate brain maps is to identify significant associations between effects expressed on the cortical mantle – where a ground truth is lacking – such as the correlation between the spatial pattern of a gene’s expression and spatially distributed activation patterns or cortical morphology ^3,5,51^. We next explored associations between the first principal component (PC1) of gene expression ^52^ (Fig. 4A, left) with well-validated surface maps (Fig. 4A, middle) of function (the principal gradient of cognitive terms from functional activation studies; *Neurosynth*) ^53,54^; structure (the average ratio of T1-weighted to T2-weighted MRI) ^47,55^; morphology (average cortical thickness) ^47,55^; and intrinsic functional connectivity (the first principal component of resting-state functional connectivity) ^56^. We performed inference on these associations using surrogates derived from eigenstrapped surrogates (blue) and compared the results to the BrainSMASH (green) and Spin Test (yellow) methods. Empirical correlations were *z*-scored to quantify the relative effect size and statistical significance of each null (Fig. 4B).

**Fig. 4.**
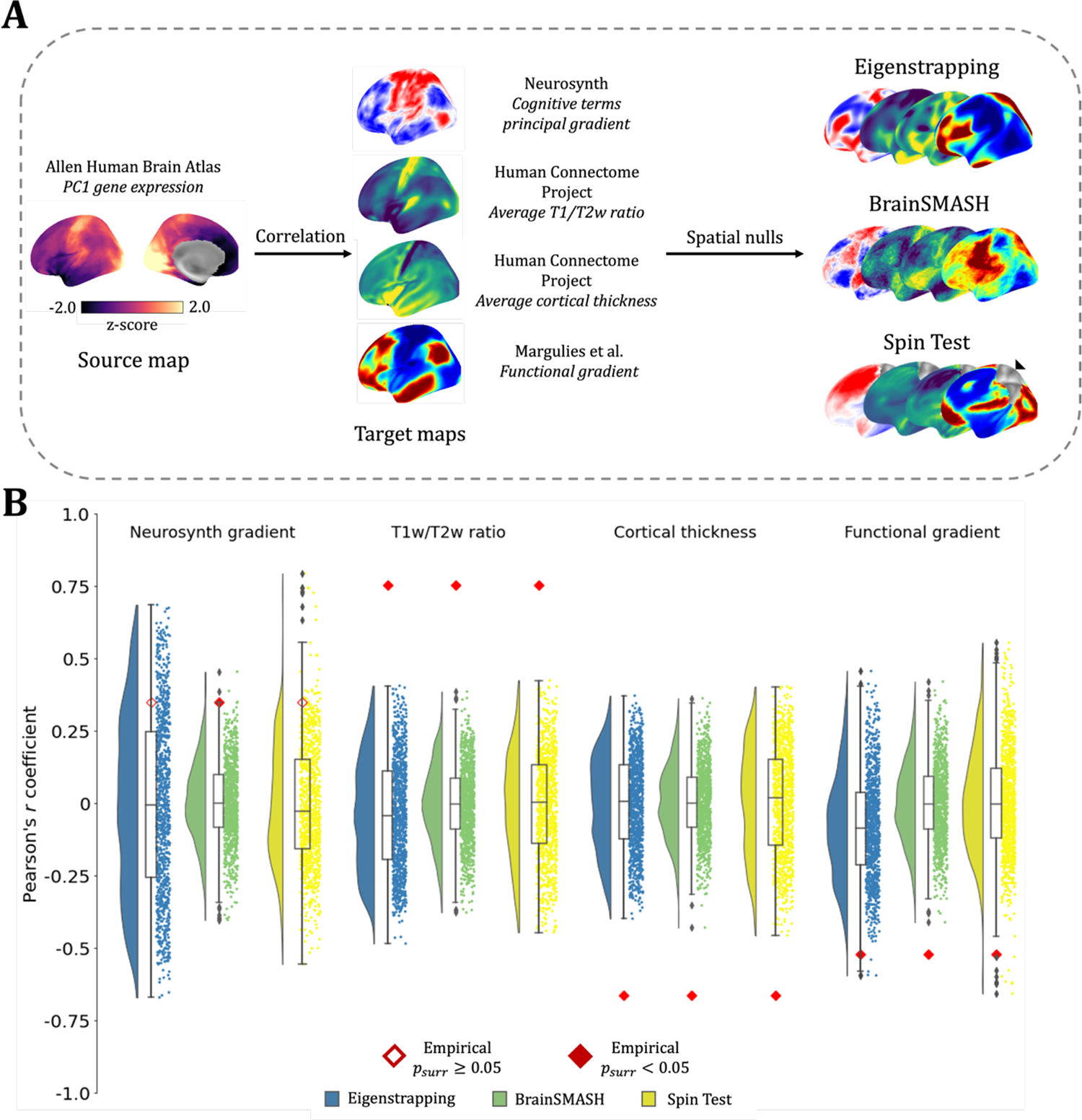
Null hypothesis testing of associations between empirical brain maps. (A) Examining the association of the first principal gradient (PC1) of cortical gene expression (left) with four example target maps (center). The exemplar surrogates per method are plotted on the left hemisphere of the inflated *fsaverage* surface. Inference on these associations was performed using three SA-preserving surrogate methods (eigenstrapping: blue; BrainSMASH: green; Spin Test: yellow) (B) Correlations with source brain map (first principal gradient of gene expression) of target brain maps (*Neurosynth gradient, T1w/T2w ratio, Cortical thickness,* and *Functional gradient*) in red; surrogate correlations to source map plotted with rainclouds ^57^ (eigenstrapping: blue; BrainSMASH: green; Spin Test: yellow). Empirical correlations of source/target pairs are given by red-bordered (non-significant, *p*_*surr*_ ≥ 0.05) or red-filled (significant, *p*_*surr*_ < 0.05) diamonds. All *p*-values are family-wise error corrected ^58^.

The correlations of each of these maps to gene expression vary considerably in magnitude and sign (see red diamonds in Fig. 4B). The association of the cognitive gradient to the gene expression map is the weakest (Fig. 4B, left; *Neurosynth gradient*). Notably, the null is only rejected for the BrainSMASH test (*z* = 2.55, *p* = 0.008) whereas the nulls derived from the two other methods possess wider tails which enclose the empirical correlation (eigenstrapping *z* = 1.14, *p* = 0.39; Spin test *z* = 1.71; p = 0.062). All other associations are statistically significant (i.e., *p* < 0.05) when using any of the nulls, although the correlation distributions are consistently narrowest for the BrainSMASH test, yielding larger *z*-statistics for the T1w/T2w ratio (eigenstrapping *z* = 4.18, BrainSMASH *z* = 5.69, Spin Test *z* = 4.18), cortical thickness (eigenstrapping *z* = −4.10, BrainSMASH *z* = −5.34, Spin Test *z* = −3.59), and the functional gradient (eigenstrapping *z* = −2.42; BrainSMASH *z* = −3.86, Spin Test *z* = −2.90).

The narrower tails of the BrainSMASH method are notable and could be due to a whitening effect on the SA, which is evident in the noisier visual appearance of these nulls (Fig. 4A, middle; Fig. S7A). Although the SA is preserved to the width of the kernel, a lack of smoothness is present at larger separation distances (i.e., the variogram is flatter; Fig. S7B). This issue does not arise with the eigenstrapped nulls (Fig. S7C). Very long-wavelength SA (captured by the eigenspectrum) is preserved with eigenstrapping but degraded by the BrainSMASH test (Fig. S7D-E). Although the Spin Test preserves the SA, the rotation of the medial wall is evident – the black marker on the Spin Test surrogate (Fig. 4A: *Spin-permuted*, rightmost brain map) indicates the non-data (NaNs) from the medial wall that are rotated onto the cortical surface. This issue is avoided by eigenstrapping, as eigenmodes are rotated in the eigenspace *Z*, rather than the cortical surface *x*.

### Generating subcortical surrogate maps

Characterizing subcortical activity and cortical-subcortical interactions is of substantial current interest ^51,56,59–64^. We extended eigenstrapping to volumetric data to enable significance testing of associations between and within subcortical structures. As a demonstration, we constructed tetrahedral meshes of three subcortical structures (thalamus, hippocampus, and striatum; see Methods) and applied eigenstrapping to these discretized surfaces. The process for generation of eigenstrapping surrogate maps in subcortical and volumetric spaces is identical to the process for cortical surfaces once a mesh has been derived. In brief, subcortical (tetrahedral) geometric eigenmodes are transformed to the spherical representation and randomly rotated, then transformed back, producing subcortical surrogate maps with matched SA as in Eq. (2). As an example, we derived maps of cortico-subcortical associations (known as “functional connectivity gradients”, see Supplementary Information-S6 for details), which capture the principal variations of functional connectivity between subcortical and cortical voxels. For present purposes, this method yields smoothly varying patterns projected onto thalamus, hippocampus, and striatum, from which we derive eigenstrapped surrogates (Fig. 5A).

**Fig. 5.**
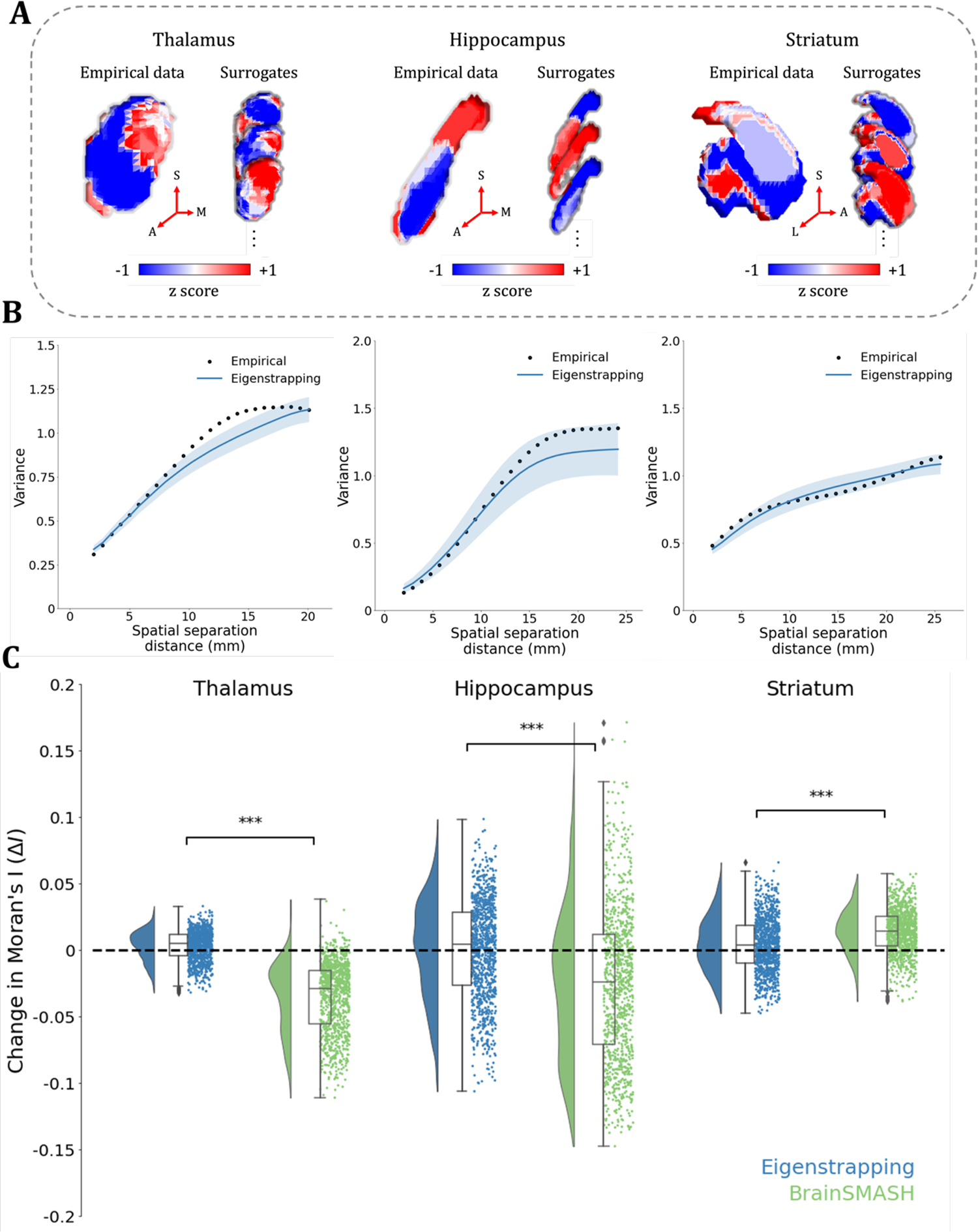
Subcortical surrogate maps with spatial autocorrelation. (A) Cortico-subcortical connectivity gradients (*empirical data*) in left thalamus (left), hippocampus (middle), and striatum (right) and 3 example surrogates generated using eigenstrapping. Number of modes used were 700, 100, and 300 for thalamus, hippocampus, and striatum, respectively, and all surrogates had amplitude adjustment applied. Thalamus surrogates also had residuals permuted. Axes of subcortical projections are given by red arrows: S: superior; A: anterior; M: medial; L: lateral. (B) Variograms of subcortical principal gradients (black) and 1000 surrogates (blue) across the three subcortical structures. (C) Change in Moran’s *I* (Δ*I*) for difference in SA within principal gradients. Rainclouds of 1000 surrogates of eigenstrapping (blue) shown against BrainSMASH (green) for each subcortical structure. Black bars denote *T*-tests performed between each null method Δ*I*. Stars correspond to significance level of two-sided *p*-values of *T*-tests: ***: *p* < 0.005; **: *p* < 0.01; *: *p* < 0.05; n.s.: *p* ≥ 0.05.

The application of eigenstrapping to these structures generates subcortical surrogates that preserve the variety of SA in these data (Fig. 5B). Eigenstrapping preserves empirical SA (change in Moran’s *I*; Δ*I*) more accurately than BrainSMASH with optimized parameters (Fig. 5C). Specifically, the changes in Moran’s *I* was significantly lower in eigenstrapping surrogates compared with BrainSMASH surrogates across all subcortical structures (*thalamus*: Student’s *T-*statistic (*T*) = 42.24, *p* < 0.0005, *degrees of freedom* (*d.f.*) =1998; *hippocampus*: *T* = 12.27, *p* < 0.0005, *d.f.* = 1998; *striatum*: *T* = −11.46, *p* < 0.0005, *d.f.* = 1998). Note that the Spin Test cannot currently generate surrogates of volumetric maps, so it could not be compared with the eigenstrapping and BrainSMASH results in these subcortical structures.

### Higher-order spatial correlations and complex textural features

While SA captures the linear, two-point smoothness of a pattern, many spatial maps possess higher order correlations, with ternary (three-point) and quaternary (four-point) relationships that cannot be predicted from knowledge of standard (two-point) correlations. These complex textures arise in systems showing accumulative and thermochemical processes such as soils ^65^, alloys ^66^, and gene enrichment in plants ^67^. Ternary and quaternary effects are also present in natural scenes ^68^, where they are central to human visual perception ^18,19,21^ and associated responses in visual cortex ^20^.

Many effects expressed on the cortex arise from complex biophysical processes. It is hence possible that many cortical maps, such as ocular dominance stripes, possess complex textural properties. Establishing their presence requires a surrogate method that preserves low order (binary) correlations but randomizes higher-order (ternary, quaternary, *etc.*) correlations. As a proof of principle, we project a human face, a canonical multiscale natural scene, to the cortex (Fig. 6A). The Spin Test rotates this cortical map and distorts but does not disrupt the complex textural relationships when projected back to the grid (Fig. 6A, bottom). It thus provides an insufficiently deep randomization of the original map. In contrast, eigenstrapping preserves two-point correlations (first-order smoothness) but visually randomizes these more complex cross-scale properties (Fig. 6A, top) due to the randomization of effects across independently rotated scales (eigengroups).

**Fig. 6.**
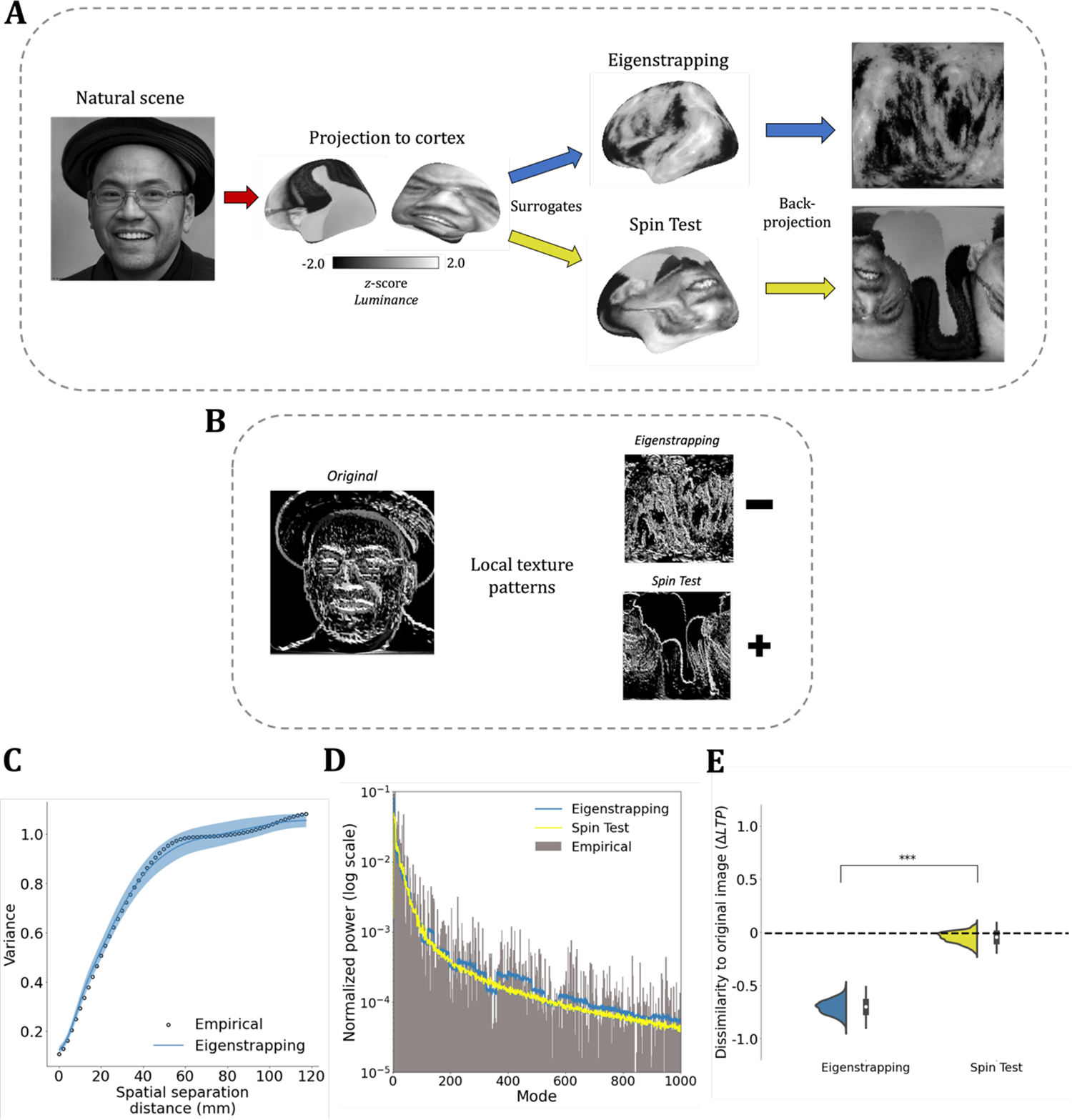
Eigenstrapping randomizes complex textural features in a natural scene. (A) A grayscale image of a natural scene (an artificially generated face) projected to the cortex. 1000 surrogates generated from eigenstrapping (blue) or the Spin Test (yellow). Luminance values are *z*-scored and kept constant throughout the analysis. Surrogates are then projected back to the square grid. (B) Grid-projected images are discretized using local texture patterns (local ternary patterns; LTP ^69^), which classify values (−1: black, 0: gray, 1: white) based on the similarity of a local neighborhood to a central pixel. (C) The variogram of the eigenstrapping surrogates follows the empirical curve from very fine to coarse spatial scales. (D) The average modal power spectra of the surrogates are nearly identical (Pearson’s *r* = 0.949) and reproduce the empirical power spectrum (gray) (Pearson’s *r* = 0.62 and 0.50 for eigenstrapping and Spin Test, respectively). (E) The proportion change in local ternary patterns 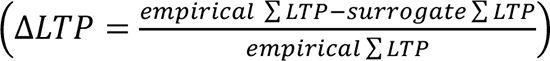 for each surrogate method.

To test this effect more formally, we discretized the images using local texture patterns (local ternary patterns; LTP ^69^), which classify values (−1: black, 0: gray, 1: white) based on the similarity of a local neighborhood to a central pixel (Fig. 6B, see Methods). Both eigenstrapping and the Spin Test preserve the variogram and the eigenspectrum of the face (Fig. 6C-D) but only eigenstrapped surrogates disrupt these textural properties (Fig. 6E). The difference between the methods’ proportion ΔLTP is substantial (T = 237.81, p < 0.0005, d.f. = 1998). Eigenstrapping thus presents a unique method to generate a null distribution for identifying complex textural properties in brain maps.

## DISCUSSION

We present a method to generate surrogate brain maps by resampling geometric basis sets. Eigenstrapping can yield a very large number of surrogate data realizations for even small surfaces such as subcortical structures, while closely preserving the spatial smoothness of the original data. These additional realizations explore a deeper null space than other methods, generating surrogate maps that preserve two-point correlations but randomize more complex textural properties. Unlike the Spin Test, eigenstrapping avoids the “medial wall problem” (see Supplementary Information-S7) and can be extended to subcortical structures. In comparison with BrainSMASH, eigenstrapping preserves the full spatial power spectrum, preserving spatial correlations well beyond the spatial smoothing kernel that lies at the core of the BrainSMASH method. For this reason, eigenstrapping preserves the Moran’s local *I* statistic more faithfully than the BrainSMASH test and does not require parametric assumptions or extensive parameter tuning, with eigenstrapping only having one free parameter (the number of modes used for decomposition in Eq. 1). Improvements over the current state-of-the-art are summarized in Table 1. We provide an open-access Python package, which implements the method for both surface and volumetric maps ^70^.

**Table 1.**
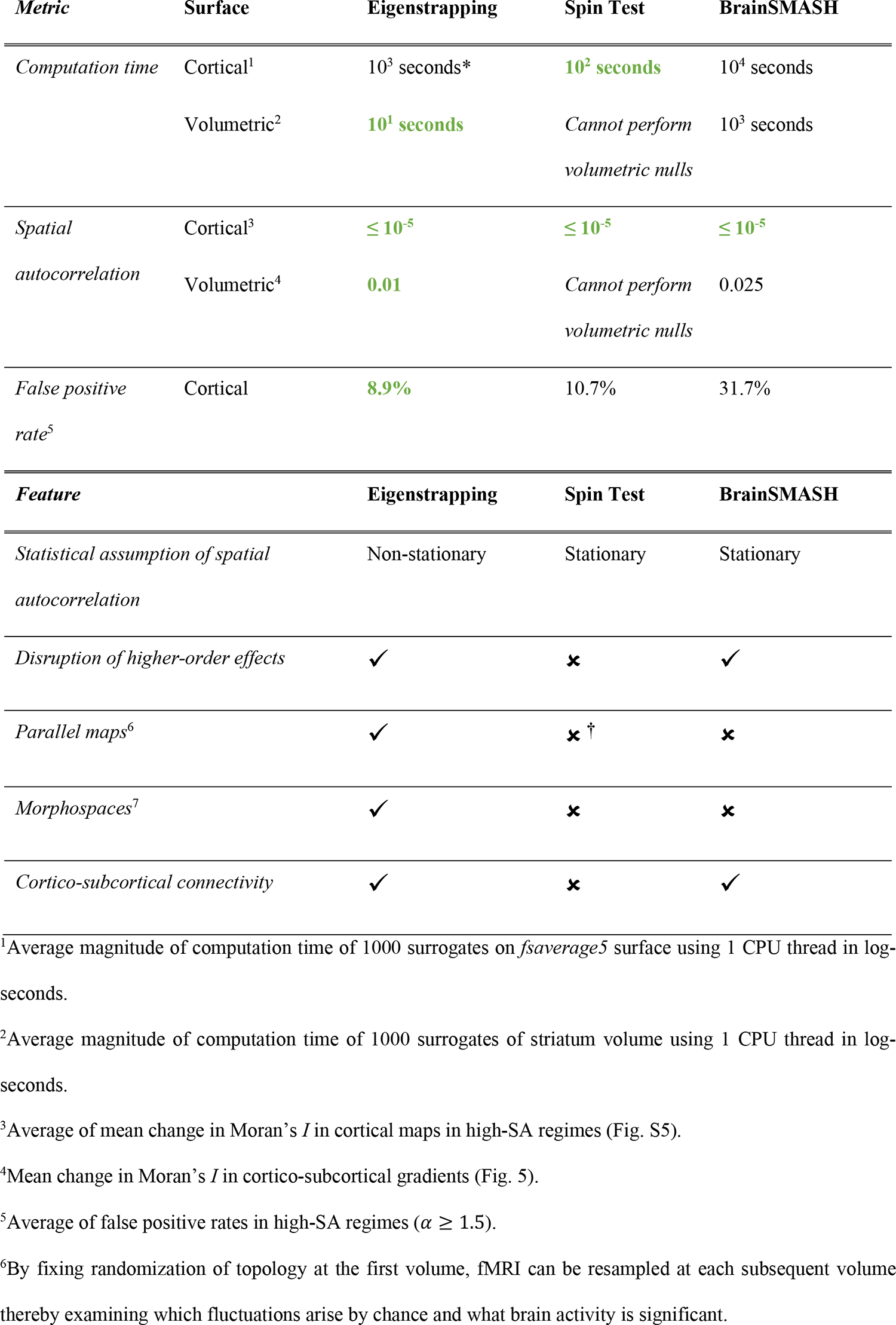
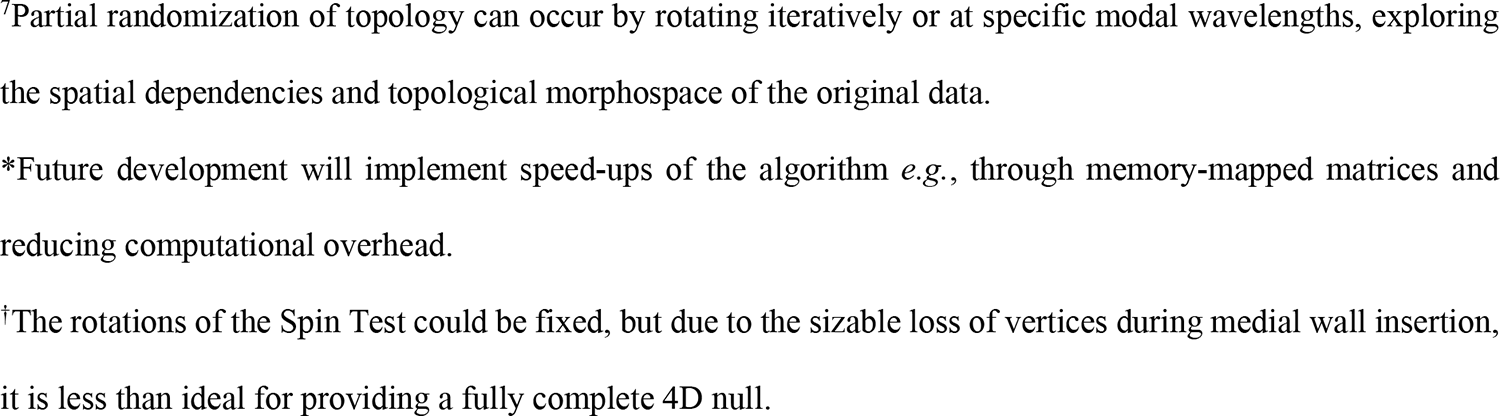
Comparison of eigenstrapping to other null models.

| *Metric* | *Surface* | *Eigenstrapping* | *Spin Test* | *BrainSMASH* |
| --- | --- | --- | --- | --- |
| *Computation time* | Cortical<sup>1</sup> | 10<sup>3</sup> seconds* | **10<sup>2</sup> seconds** | 10<sup>4</sup> seconds |
|  | Volumetric<sup>2</sup> | **10<sup>1</sup> seconds** | Cannot perform<br>volumetric nulls | 10<sup>3</sup> seconds |
| *Spatial autocorrelation* | Cortical<sup>3</sup> | **≤ 10<sup>-5</sup>** | **≤ 10<sup>-5</sup>** | **≤ 10<sup>-5</sup>** |
|  | Volumetric<sup>4</sup> | **0.01** | Cannot perform<br>volumetric nulls | 0.025 |
| *False positive rate<sup>5</sup>* | Cortical | **8.9%** | 10.7% | 31.7% |
| *Feature* |  | *Eigenstrapping* | *Spin Test* | *BrainSMASH* |
| *Statistical assumption of spatial autocorrelation* |  | Non-stationary | Stationary | Stationary |
| *Disruption of higher-order effects* |  | ✓ | ✗ | ✓ |
| *Parallel maps<sup>6</sup>* |  | ✓ | ✗<sup>†</sup> | ✗ |
| *Morphospaces<sup>7</sup>* |  | ✓ | ✗ | ✗ |
| *Cortico-subcortical connectivity* |  | ✓ | ✗ | ✓ |

<sup>1</sup>Average magnitude of computation time of 1000 surrogates on *fsaverage5* surface using 1 CPU thread in log-seconds.
<sup>2</sup>Average magnitude of computation time of 1000 surrogates of striatum volume using 1 CPU thread in log-seconds.
<sup>3</sup>Average of mean change in Moran's *I* in cortical maps in high-SA regimes (Fig. S5).
<sup>4</sup>Mean change in Moran's *I* in cortico-subcortical gradients (Fig. 5).
<sup>5</sup>Average of false positive rates in high-SA regimes ( $\alpha \geq 1.5$ ).
<sup>6</sup>By fixing randomization of topology at the first volume, fMRI can be resampled at each subsequent volume thereby examining which fluctuations arise by chance and what brain activity is significant.
<sup>7</sup>Partial randomization of topology can occur by rotating iteratively or at specific modal wavelengths, exploring the spatial dependencies and topological morphospace of the original data. \*Future development will implement speed-ups of the algorithm *e.g.*, through memory-mapped matrices and reducing computational overhead. <sup>†</sup>The rotations of the Spin Test could be fixed, but due to the sizable loss of vertices during medial wall insertion, it is less than ideal for providing a fully complete 4D null.

By rotating spatial modes within groups and re-inserting the coefficients from the original eigenmode decomposition, eigenstrapping preserves the average amplitude of each spatial frequency, and as such is a natural extension of time series phase randomization ^41^ and wavestrapping ^71,72^, to spatial data on curved and folded surfaces. We show that eigenstrapping permutes higher order (ternary) properties of spatial maps, just as phase randomization permutes comparable nonlinear properties in time-series data. Though it is generally accepted that the brain expresses nonlinear activity ^24–27,73^, the presence and putative function of nonlinear properties of brain maps is an empirical question for which we provide the inferential tools.

Eigenstrapping lends itself to an extension to spatiotemporal data, again by importing a technique from multivariate phase randomization of time series data ^71,74^: Applying the same random rotation of each eigengroup across whole brain volumes acquired sequentially through time preserves temporal properties of each point-wise timeseries and spatial relationships between timeseries, while randomizing all other spatial properties. Excursions outside this spatiotemporal null would be informative regarding complex physiological processes, such as the presence of travelling waves ^28,31,32,36,39,75^ and metastable dynamics ^25,26,30^. More broadly, any metric sensitive to time-dependent functional connectivity could be employed to detect non-trivial fluctuations in brain state, which are of substantial current interest ^76–83^. Demonstrating these effects will be the subject of future work.

Identifying a suitable orthogonal transformation that removes the complex correlations within spatiotemporal data is key to nonparametric methods, as this allows rotation of the phases of basis functions without degrading the correlations of the original data ^84,85^. Resampling methods for null hypothesis testing of neuroimaging data have previously employed the discrete wavelet transform for this endeavor ^72,86^. However, while wavelet-based resampling methods are suitable for data on regular two-dimensional grids (such as fMRI slices) ^72^, the geometric distortions induced by cortical curvature place limitations on the application of wavelet-based methods to contemporary surface-based analyses ^59^. Obtaining geometric eigenmodes from the LBO is a natural extension of orthogonal basis decompositions to curved surfaces and, as shown here, yield surrogate brain maps that preserve SA and provide good control of false positives. A related approach, known as Moran spectral randomization (MSR), first weights vertex connectivity of the surface (usually using the inverse of the pairwise distance matrix) and then estimates the graph Laplacian of this matrix. The ensuing eigenfunctions of the generalized eigenvalue problem for the Laplacian are then used to decompose a map on the surface (similar to Eq. 1). Surrogate maps are derived by randomly flipping the (positive or negative) sign of the coefficients ^87^. This process yields 2^*n*–1^ surrogates, where *n* is the total number of eigenfunctions, far fewer than arising from free rotation of geometric eigengroups as in this paper, which is (*G* − 1)!. Moreover, if the spatial map loads onto a small number of the eigenfunctions, as often happens, flipping the sign of the coefficient yields surrogates that are strongly (anti-)correlated to the original data, producing multimodal null distributions ^8^. As a result, the MSR achieves poorer FPR in similar tests ^11^.

Geometric eigenmodes and their associated eigenvalues are obtained by solving the Helmholtz equation on a discrete cortical mesh (see Methods). As such, geometric eigenmodes play a crucial role in generative models of brain activity ^30,33,34^ and morphology ^88^. In particular, physiologically derived neural field models ^89^ are separable into their temporal and spatial components under very broad assumptions ^33,34^. The spatial component of a broad class of neural field models satisfies the Helmholtz equation, yielding the geometric modes that we presently employ. These modes thus capture how geometry constrains large-scale neural activity ^43^. The temporal component of neural field models assigns damped oscillations to each eigenmode (higher frequencies are associated with eigenmodes with shorter characteristic wavelengths). Although we use geometric eigenmodes for a specific statistical purpose, this deeper connection to neural field theory (NFT) could assist in linking the statistical inference that eigenstrapping affords to deeper causal inference. For example, finding a significant excursion of cortical activity from an eigenstrapped null could motivate exploration of linear resonance or nonlinear excitation of NFT, extending prior work from a purely temporal to a spatiotemporal framework ^90,91^.

Eigenstrapping is a very versatile approach, being applicable to both surface- and volume-based analyses. It is also fast for most applications, with 200–500 modes being adequate to randomize common neuroimaging datasets, such as smoothed fMRI maps, while preserving intrinsic spatial structure. We also note that incremental rotation of eigengroups (applying a series of random but small rotations) would allow one to track the gradual randomization of a brain map through a complex morphospace ^92^, similar to the approach recently applied to synthetic brain networks ^93^ and natural images ^20^. In sum, eigenstrapping offers a flexible methodology for null hypothesis testing in modern neuroscience.

## METHODS

### Derivation of geometric eigenmodes

Cortical eigenmodes were derived from a triangular mesh of a population-averaged template left hemisphere pial surface with vertices *x* ^94^ (*fs-LR-32k* template space with 32,492 vertices per hemisphere ^55,95^: Fig. 1A, gray cortical surface. *fsaverage5* template space with 10,242 vertices per hemisphere ^94^). The eigenmodes and associated eigenvalues are obtained by solving the Helmholtz equation,

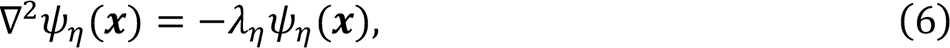

where ∇^2^ is the Laplace Beltrami Operator (LBO) which generalizes the Laplacian to the curved surface ^94,96^ where η = 0,1,2, … indices the eigenmodes. The geometric modes ψ_η=Λμ_ = Sψ_0,0_(*x*), ψ_1,–1_(*x*), ψ_1,0_(*x*), … ψ_Λ,μ_(*x*)T have corresponding eigenvalues λ_η=Λμ_ = Sλ_0,0_, λ_1,–1_, λ_1,0_ …, λ_Λ,μ_(*x*)T, where the index of mode η becomes group Λ and number μ. When the Helmholtz equation is applied to study waves, the eigenvalues are typically denoted by λ = *k*^2^ where *k* is known as the wave number ^33^. These groups increase in size monotonically according to the multiplicity factor *n* = 2Λ + 1 and decrease in spatial wavelength with group (Fig. S10). The integer μ in each group Λ ranges from −Λ ≤ 0 ≤ Λ. The eigenmodes ψ_η_(*x*) form a complete set of orthonormal basis functions, hence supporting the decomposition of a surface map into space-varying components with coefficients β (as in Eq. 1).

The Laplace-Beltrami operator (LBO) is a generalization of the Laplacian on a sphere ***z*** to functions defined on arbitrary smooth surfaces such as the cortex ***x***. On the sphere, the solutions to Eq. (6) are identical to spherical harmonics, which are denoted *Y*_*lm*_(*Z*) with eigenvalues ξ_*lm*_.

Note that spherical harmonics occur in groups *l* which are rotationally invariant, hence with identical eigenvalues ξ_*lm*_ = ξ_*ln*_ where *n* ≠ *m* index these degenerate, within-group harmonics. In the non-spherical case (e.g., on the cortex), eigenvalues λ_Λμ_ ≠ λ_Λ/_ where *v* ≠ μ and are no longer degenerate (Fig. S1).

Perturbation theory can be used to express LBO eigenfunctions ψ_*lm*_and eigenvalues λ_*lm*_on non-spherical surfaces in terms of first order perturbations of the spherical harmonics ^34^. For the geometric eigenvalues from Eq. (5) this corresponds to

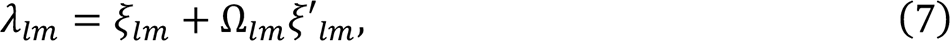

where ξ_*lm*_is the unperturbed (spherical harmonic) eigenvalue and ξ′_*lm*_is the first order perturbation with coefficient Ω_*lm*_. Hence, this expresses eigenmodes and their eigenvalues as a first order perturbation of spherical harmonics. We use the indexing η = Λμ with group Λ and mode number μ in the geometric (cortical and subcortical) case, and η = *lm* in the spherical case to disambiguate geometric eigenmodes and spherical harmonics.

Crucially, the geometric eigenvalues within groups are perturbed by differing amounts because of the symmetry breaking transformation of the sphere onto the folded cortex. That is, the perturbation Ω_*lm*_ ≠ Ω_*ln*_ and thus the eigenvalues λ_*lm*_ ≠ λ_*ln*_ when *m* ≠ *n*. Fig. S1 shows the first 16 eigenvalues obtained by solving Eq. (5) on increasingly folded cortices from a spherical representation (far left; ρ = 0) to a fully folded cortex (far right; ρ = 1) using FreeSurfer ^94^, as a function of folding ρ. As ρ increases, the average eigenvalue within groups Λ remains nearly constant (exactly so for the zeroth group Λ = 0) while the eigenvalues for individual modes demonstrate perturbed energies, splitting but not crossing with modes from adjacent groups.

Eigenstrapping proceeds by rotating these geometric eigenmodes collectively within groups, hence maintaining within-group orthogonality while achieving rotations across spatial scales (perturbing higher-order patterns, in contrast to the Spin Test). To achieve this without distorting the associated eigenvalue energies of different modes within a group, we restore the symmetry of eigenvalues within groups through renormalization. This renormalization can be recast as a mapping of geometric modes on the surface of a hypothetical ellipsoid with dimension *n* = 2Λ + 1 and axes λ_Λμ_ to spherical modes on an n-dimensional (hyper)sphere with axes ξ_*lm*_(Eq. 2). That is, the first non-constant spherical harmonic (and modal) group forms the surface of a 3D sphere, while the second group forms the surface of a 5D sphere, and so on.

The composition of these eigengroups and the geometric properties of each ellipsoid are crucial to the eigenstrapping approach. The approximate wavelength on the cortex can be calculated for each cortical eigengroup as ^43,88^

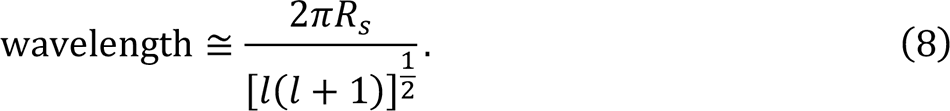

The wavelengths for a sphere of radius *R*_s_ ≈ 67 mm (approximately the radius of the *fsaverage5* population-average template used in this study) are listed in Supplementary Table 1, along with eigengroup membership for the first 1000 modes. The linear relationship of eigengroup membership and the relationship of wavelength to group is given in Fig. S10. Fig. S10 also shows the group size for the first 100 eigengroups, corresponding to the first 10000 modes.

Once transformed to an *n*-dimensional sphere with discrete grid points *Z*, each spherical mode Ø_Λμ_(*Z*) can be expressed as a weighted sum of spherical harmonics *Y*_*lm*_(*Z*),

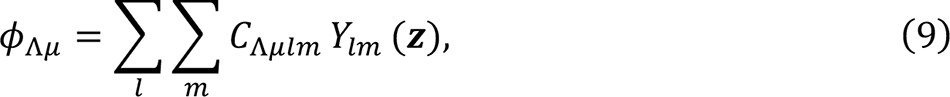

where *C*_Λμ*lm*_ are coefficients of this expansion of spherical eigenmodes Ø_Λμ_(*Z*) in terms of spherical harmonics *Y*_*lm*_(*Z*). For low order modes, the terms on the RHS are predominated by harmonics from the same group as the spherical eigenmodes, that is *C*_Λμ_ ≫ *C*_*lm*_ for groups *l* ≠ Λ ^33^. For higher modes, there is greater “leakage” from adjacent groups ^97^. Note that geometric eigenmodes can be more complex than their corresponding spherical harmonics, such as differing numbers of positive and negative domains within the same group. However, the Courant Nodal Line Theorem limits the complexity of the resulting modes by restricting the number of separate regions that can have positive or negative sign to at most *n* for the *n*th eigenmode ^33^.

Having mapped the modes to the sphere, the ensuing spherical eigenmodes Ø_Λμ_ can be rotated in blocks of size *n* = 2Λ + 1 to yield new modes with the same characteristic wavelength. Rotating all modes within each block by a common angle ensures they remain orthogonal to each other.

### Rotation of spherical modes

Rotation of spherical modes φ is the matrix multiplication of mode groups of size {*Z* × *n*}, where *Z* is the number of vertices in the abstract (hyper-)sphere *Z* and *n* is the number of modes in that group, by a random rotation matrix *R*(θ_Λ_) ^98,99^ of size {*n* × *n*},

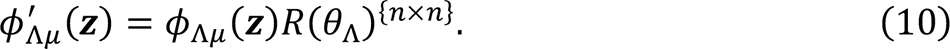

where θ_Λ_ is the random angle, drawn randomly for the Λ-th group. Eq. (10) is performed on all groups, resulting in unbiased rotations of modes that preserves eigenvalue energies and within- and between-group orthogonality. To generate random rotation matrices on the sphere *Z*, we draw from the Haar distribution for the special orthogonal group SO(*n*) ^98^, where *n* is the number of modes in the group Λ. This is performed in practice by taking random selections of SO(*n*) using the *scipy.stats.special_ortho_group* Python function ^99^.

### Generalized linear model for generating surrogate maps

An empirical brain map *y*(*x*) on spatial location *x* with components {*x*, *y*, *z*} can be decomposed as a weighted sum of geometric modes ψ_Λμ_(Eq. 1). The weightings, or coefficients, β_Λμ_ of this sum can be obtained by integrating over the cortical surface ^34,44,45^,

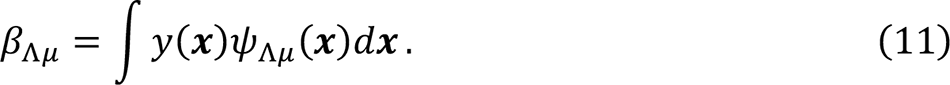

Average normalized modal coefficients 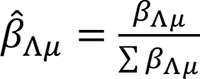 for HCP data (used in Fig. 1) are plotted in Fig. S11.

### Eigenstrapping algorithm

The algorithm for rotating eigengroups and creating randomized, SA-preserving surrogate brain maps (the eigenstrapping algorithm; Algorithm 1) can be outlined in pseudocode, as follows:

**Algorithm 1.**
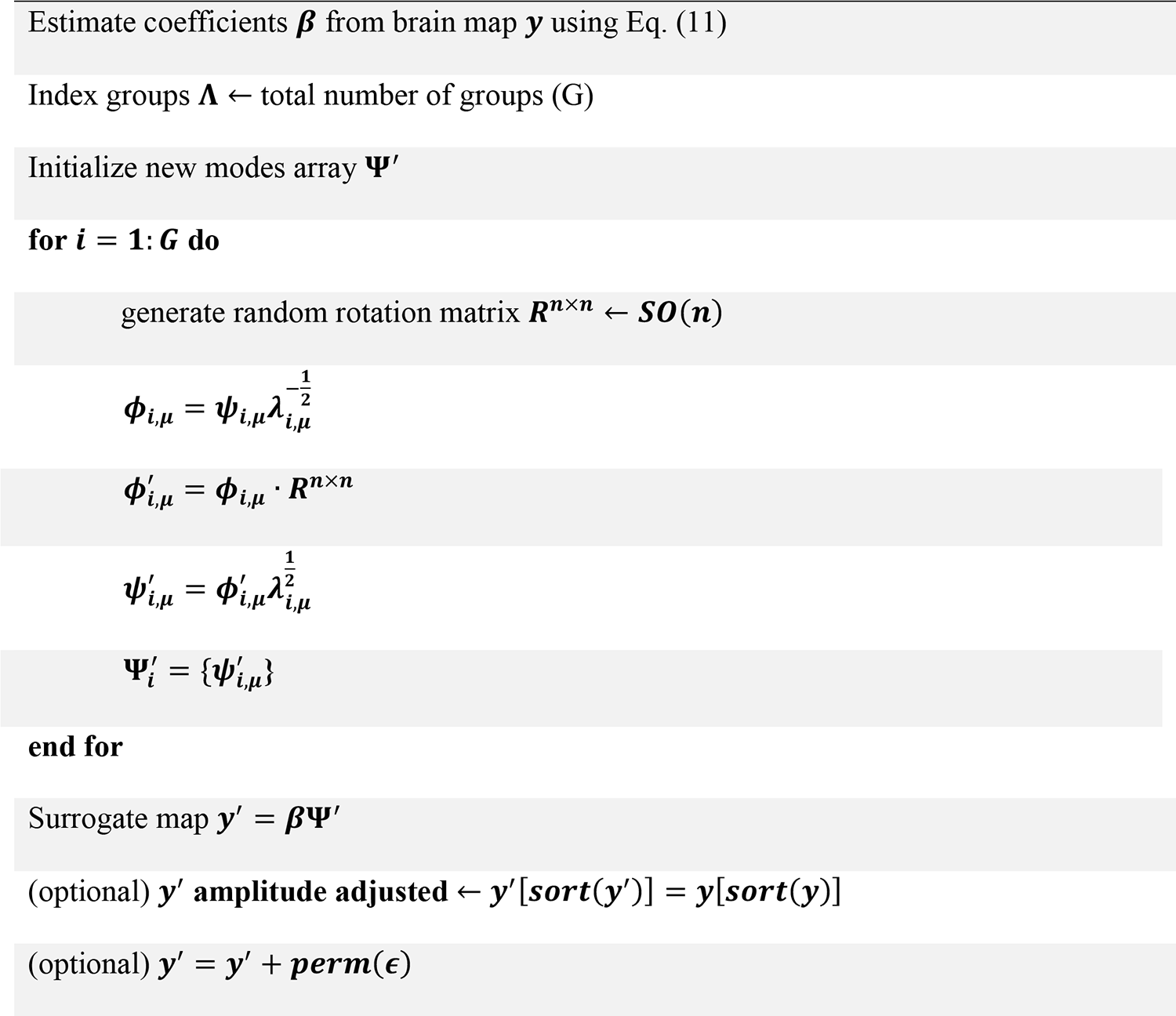
Eigenstrapping algorithm

Note that the operator (′) refers to ‘new’, or ‘prime’, not the first derivative or the transpose.

Eigenvalues λ_Λμ_ and eigenmodes ψ_Λμ_ were derived from the left *fs-LR-32k* and *fsaverage5* pial surfaces using a numerical finite element method and the Lanzcos algorithm of the Implicitly Restarted Arnoldi Method in ARPACK ^100^ as implemented in the ShapeDNA Python package ^101,102^ (https://github.com/Deep-MI/LaPy). These surfaces consist of 32,492 and 10,242 vertices, respectively. For Figs. 1-2 (on the *fs-LR-32k* pial surface), we computed up to the first 6,000 modes to test the algorithm, which corresponds to approximately 18.5% of the complete set of surface modes. For Fig. 3 in the simulated maps (on the *fsaverage5* pial surface), we computed up to the first 2500 modes, corresponding to approximately 2.5% of all surface modes. In simulated maps that resembled brain data (1.5 ≥ α ≥ 3.0), the first 200 to 1000 modes were adequate to produce surrogate maps with the same SA structure (corresponding to 0.2 to 1.0% of all surface modes). All cortical surfaces had the medial wall removed (cut and turned into a Neumann boundary) prior to deriving eigenvalues and modes, except as addressed in the texture analysis section (Methods: Local texture patterns; Fig. 6). The Neumann boundary forces the derivative at the normal to the boundary to zero. This boundary condition was chosen rather than the Dirichlet condition due to its better minimization of boundary effects ^101,102^.

There are several advantages of this method:

1. Because modes are generalizable representations of real-valued functions on the surface, they can produce surrogate brain maps from any surface that is utilized in imaging analyses (e.g., cortical maps across the entire developmental spectrum as well as from different species).
2. Volumetric modes can be calculated using tetrahedral meshes that represent volumetric space and projected back to volume space. Thus, null maps can also be derived for volume maps as well as surface maps. This is important as subcortical spatiotemporal maps are a topic of substantial growing interest ^59,61,62,64,103^.
3. As boundary effects are minimized ^101^, modes can be derived without the presence of the medial wall, as they were in all cortical analyses in this paper, or for an arbitrarily-sized patch of brain for region-of-interest analysis, such as insular or hippocampal maps ^59,104^.
4. One can derive a number of rotated modes Ψ* on any finite mesh to draw from and create practically infinitely many different surrogate maps at will. The computation time of Alg. (1) was calculated for different surfaces and total numbers of modes in Fig. S9.
5. In contrast to modes derived from the graph Laplacian of the surface distance matrix (i.e., for MSR ^87^), geometric eigenmodes can be smoothly rotated. This permits a far larger space of nulls than MSR and avoids producing nulls that are strongly (anti-)correlated with the original data ^8^.

### Depth of decomposition and treatment of residuals

The eigenmode decomposition Eq. (1) is complete for a total of *N* modes when performed on a spatial mesh with *N* vertices. Accordingly, the residual term ε_*G*_(*x*) in Eq. (1) approaches zero (within numerical accuracy). However, for a highly resolved cortical mesh (e.g., 32,492 vertex points for *fs-LR-32k*), a complete representation carries a substantial computational burden. In practice, the error term becomes small for substantially fewer eigenmodes than the above limit. Retaining the variance of the error ε is important to preserve the power and variogram of the original data (Fig. S2A). However, simply adding the original error yields surrogate data that is correlated with the original data (Fig. S2B-C) when the number of modes is not sufficiently high enough. This is because exactly the same error term ε(*x*) is present in all surrogate realizations. Randomly permuting the residual to yield a surrogate residual ε* ensures the correlation between data and surrogate (and between surrogates) is zero-centered. However, if the error term is too large, this process will whiten the data, distorting the variogram (Fig. S3). In practice, we found that selection of eigenmodes sufficient to balance parsimony with accuracy (i.e., preserving the variogram and ensuring a zero centered correlation between *y* and *y’*) is an important parameter to optimize – ranging from 50–2000 in the GRF analysis (Fig. 2) to 6000 in the HCP data (Fig. 1). For the smaller subcortical meshes, between 100–700 eigenmodes were sufficient (Fig. 5). As a rule of thumb, it is recommended that 200–500 modes be sufficient for most smoothed fMRI data. With parcellated or very smooth data, fewer modes are required. Tools have been provided for the user to fit the number of modes to the empirical SA in the Python toolbox ^42^.

### HCP data

The Human Connectome Project (HCP) provides well-documented and robust activation maps for each of its task conditions ^55,105^. We downloaded activation maps in CIFTI format in *fs-LR-32k* standard space from 255 unrelated subjects for the *social cognition task*, *motor task, gambling task, working memory task, language task, emotion task,* and *relational task*, accessing 47 task contrasts in total (see Supplementary Information-S3).

### Gaussian random fields

The statistical properties of eigenstrapping were benchmarked against a known relationship using parametric simulations of Gaussian random fields (GRFs), adopting a previous approach^11^. Simulated brain maps were derived by generating pairs of 3D multivariate Gaussian distributions with Pearson correlation tuned to *r* = 0.15 ± 0.005. These ensembles of pairs were generated with nine different levels SA across by modifying the slope of the pair’s power spectral density (α = 0.0–4.0 in increments of 0.5, where 0.0 indicates random Gaussian noise). The pairs of GRFs were projected to the fsaverage5 cortical mesh using FreeSurfer *mri_vol2surf*. The medial wall was removed (*i.e.*, set to NaN), and the resulting brain maps were only included for further analysis if the intra-pair correlation fell within the target range of *r*. At each level of SA, we generated 1000 pairs, resulting in 9000 total pairs of maps with correlations from 0.145–0.155 (see Supplementary Information-S4).

### Null comparison – Spin Test

The Spin Test randomizes the alignment between two cortical surface maps through rotation by a random angle ^10^, and is useful for computing SA-corrected *p*-values when making statistical inferences on dense cortical brain maps. To compare eigenstrapping to the null brain maps generated using the Spin Test, we used the Python implementation of the method in the *neuromaps* toolbox ^5^. Any statistics drawn from Spin Test generated maps excluded the rotated medial wall (as NaNs) as the standard implementation. As the spin test can only produce null maps of the cortical surface, we used another method for comparison of both cortical surface measures and volumetric measures, namely, the BrainSMASH method, outlined below.

In contrast to the single spin per surrogate property of the Spin Test, rotations within eigenstrapping are applied group-wise to separate spherical eigenspaces, greatly increasing the true degrees of freedom. Notably, for every surrogate realized by the Spin Test, free rotation of geometric eigengroups permits (*G* − 1)! realizations. These extra realizations explore a deeper null space, where higher-order correlations are disrupted while still preserving smoothness of the original map.

### Null comparison – BrainSMASH

The Brain Surrogate Maps with Autocorrelated Spatial Heterogeneity (BrainSMASH) method uses geostatistical methods to derive randomized brain maps that replicate the empirical map’s SA ^8^. The steps that the BrainSMASH tool uses is as follows: 1) randomly permute the values in a target brain map; and 2) smooth and rescale the permuted map to recover the SA structure of the target brain map. This is performed through rescaling of values in several spatial levels of linear fits of Gaussian, exponential, or logarithmic distributions. To generate null brain maps using the BrainSMASH method, we used the Python implementation from https://github.com/murraylab/brainsmash. The dense sampling algorithm (*brainsmash.mapgen.Sampled*) was used for all analyses in this study. As the method allows for different fits to the variogram depending on parameters given, optimized parameters were chosen based on visual assessment of best fit to the original variogram.

### Cortico-subcortical functional connectivity patterns

We used resting-state functional connectivity patterns (“gradients”) to examine the capacity of eigenstrapping to identify cortical-subcortical effects. Resting-state data from the HCP were sampled on tetrahedral meshes (thalamus, hippocampus, and the striatum - consisting of the caudate, putamen, and nucleus accumbens areas). These structures were generated from binarized images of 25% probability thresholds of the Harvard-Oxford subcortical atlas of each region ^106–109^. Unlike the cortex, which can be modeled as a 2D sheet, subcortical volumes are solid 3D objects. We therefore calculated the modes using tetrahedral meshes rather than triangular meshes to account for the full 3D geometry of these structures ^102^.

Cortico-subcortical functional gradients were derived from diffusion map embedding of HCP resting-state fMRI (see Supplementary Information-S6). The first non-zero gradient for each subcortical structure (corresponding to the second eigenvector in Eq. S5) was derived from the Laplacian of the group-averaged resting state functional connectivity matrix of HCP data in MNI152 space. Eigenmodes were derived on the tetrahedral mesh, resampled to volumetric space, then rotated, yielding SA-preserving surrogate subcortical maps (Fig. 5).

### Local texture patterns

A variety of spatial images and processes, such as natural scenes, possess complex spatial features that cannot be fully captured using standard first-order statistics (two-point correlations, or spatial autocorrelation) ^68^. We refer to these higher order effects as textures to emphasize the complex arrangement of spatial features above low-level smoothness. Local texture patterns (local ternary patterns; LTPs) ^69^ were analyzed to quantify the effect of eigenstrapping on the presence of these features in natural scenes. By discretizing an image into three values (−1, 0, 1), LTPs can be used to detect complex, textural effects such as facial features.

A 1024×1024 pixel grayscale image of a face (derived from ^110^, *Natural scene*) was projected to the *fsaverage5* cortical surface (10,242 vertices) using a simple (inverse)-stereographic projection (Fig. 6A, *Projection to cortex*). The edges of the image joined at the central sulcus. Eigenstrapping (blue) with 5000 modes solved on the cortical surface (including the medial wall) was used to produce 1000 surrogates. The Spin Test (yellow) was used to rotate the data on the cortical surface 1000 times, producing 1000 rotated surrogates. The medial wall was included in this analysis in order to preserve the original luminance histogram and distortion induced by the projection. Cortical data (original image and surrogates) were then projected to the grid (*Back-projection*) by inverting the initial projection, interpolating with a bilinear spline across non-data pixels induced by projection to the grid. Central pixels *c*_*i*_ with index *i* were discretized into three values *LTP*_*i*_ = (−1,0,1) by a threshold *k* on 8-neighbor pixels *p*_*i*_

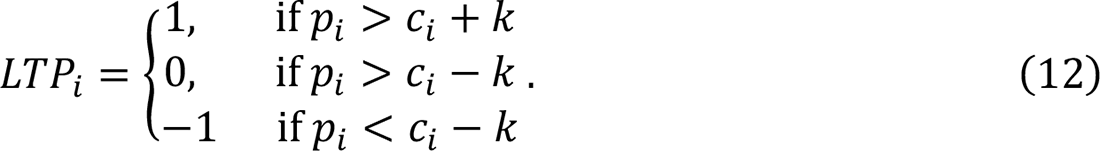

We used a threshold of *k* = 5 which is commonly used for face detection algorithms ^69,111^. Each pixel then had a value ranging from −8 to 8, which were then thresholded to positive non-zero values, indicating a common feature to a particular 9-pixel neighborhood. Each of these values were summed, producing a single LTP (∑ *LTP*) for each image. If the feature of interest (the face) is disrupted, then the LTP will be different to the original image. The proportion change in LTP (Δ*LTP*) can be quantified by the equation,

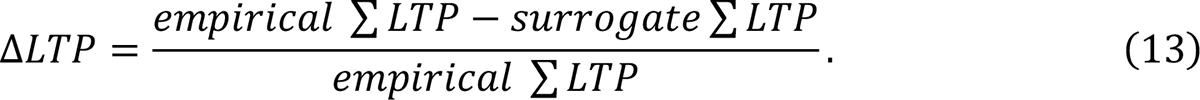

This value was derived for each surrogate and summarized in Fig. 6E. To test for a difference in the impact of the two surrogate methods’ on complex patterns, a Student’s *T*-test was performed on Δ*LTP* of each method and two-tailed *p*-values derived.

## CODE AVAILABILITY

All code and data to reproduce the results of the paper as well as the eigenstrapping method itself is openly available at https://github.com/SNG-newy/eigenstrapping/ and ^42^. Documentation on the method’s practical usage is also available from the same location. Raw and preprocessed HCP data can be accessed at https://db.humanconnectome.org/.

## ACKNOWLEDGEMENTS

We would like to thank Richa Phogat for her helpful and insightful comments on derivation of the eigenmode basis transformation. N.C.K was supported by an Australian Research Training Program Scholarship and reports no competing interests. J.C.P was supported by the Monash FMNHS Early Career Postdoctoral Fellowship and reports no competing interests. J.J reports no competing interests. B.P reports no competing interests. A.F was supported by the Australian Research Council (ID: FL220100184) and the National Health and Medical Research Council (ID: 11987431) and reports no competing interests. P.R reports no competing interests. M.B was supported by the National Health and Medical Research Council (APP1152623 and APP2008612) and reports no competing interests.

## REFERENCES

1. Hansen, J. Y. et al. Mapping neurotransmitter systems to the structural and functional organization of the human neocortex. Nat. Neurosci. 25, 1569–1581 (2022).

2. Arnatkeviciute, A., Fulcher, B. D., Bellgrove, M. A. & Fornito, A. Imaging Transcriptomics of Brain Disorders. Biol. Psychiatry Glob. Open Sci. 2, 319–331 (2022).

3. Fornito, A., Arnatkevičiūtė, A. & Fulcher, B. D. Bridging the Gap between Connectome and Transcriptome. Trends Cogn. Sci. 23, 34–50 (2019).

4. Rubinov, M. & Sporns, O. Complex network measures of brain connectivity: uses and interpretations. NeuroImage 52, 1059–1069 (2010).

5. Markello, R. D. et al. neuromaps: structural and functional interpretation of brain maps. Nat. Methods 19, 1472–1479 (2022).

6. Hansen, J. Y. et al. Mapping gene transcription and neurocognition across human neocortex. *Nat*. Hum. Behav. 5, 1240–1250 (2021).

7. Liu, H. et al. Single-cell DNA methylome and 3D multi-omic atlas of the adult mouse brain. Nature 624, 366–377 (2023).

8. Burt, J. B., Helmer, M., Shinn, M., Anticevic, A. & Murray, J. D. Generative modeling of brain maps with spatial autocorrelation. NeuroImage 220, 117038 (2020).

9. Viladomat, J., Mazumder, R., McInturff, A., McCauley, D. J. & Hastie, T. Assessing the significance of global and local correlations under spatial autocorrelation: A nonparametric approach. Biometrics 70, 409–418 (2014).

10. Alexander-Bloch, A. F. et al. On testing for spatial correspondence between maps of human brain structure and function. NeuroImage 178, 540–551 (2018).

11. Markello, R. D. & Misic, B. Comparing spatial null models for brain maps. NeuroImage 236, 118052 (2021).

12. Váša, F. & Mišić, B. Null models in network neuroscience. Nat. Rev. Neurosci. 23, 493– 504 (2022).

13. Váša, F. et al. Adolescent Tuning of Association Cortex in Human Structural Brain Networks. Cereb. Cortex 28, 281–294 (2018).

14. Kuhn, H. W. The Hungarian method for the assignment problem. Nav. Res. Logist. Q. 2, 83–97 (1955).

15. Baum, G. L. et al. Development of structure–function coupling in human brain networks during youth. Proc. Natl. Acad. Sci. 117, 771–778 (2020).

16. Cornblath, E. J. et al. Temporal sequences of brain activity at rest are constrained by white matter structure and modulated by cognitive demands. *Commun*. Biol. 3, 1–12 (2020).

17. Burt, J. B. et al. Hierarchy of transcriptomic specialization across human cortex captured by structural neuroimaging topography. Nat. Neurosci. 21, 1251–1259 (2018).

18. Tesileanu, T. et al. Efficient coding of natural scene statistics predicts discrimination thresholds for grayscale textures. eLife 9, e54347 (2020).

19. Victor, J. D., Thengone, D. J., Rizvi, S. M. & Conte, M. M. A perceptual space of local image statistics. Vision Res. 117, 117–135 (2015).

20. Puckett, A. M. et al. Manipulating the structure of natural scenes using wavelets to study the functional architecture of perceptual hierarchies in the brain. NeuroImage 221, 117173 (2020).

21. Hermundstad, A. M. et al. Variance predicts salience in central sensory processing. eLife 3, e03722 (2014).

22. Heitmann, S. & Breakspear, M. Putting the “dynamic” back into dynamic functional connectivity. Netw. Neurosci. 02, 150–174 (2018).

23. van den Heuvel, M. P., Kahn, R. S., Goñi, J. & Sporns, O. High-cost, high-capacity backbone for global brain communication. Proc. Natl. Acad. Sci. U. S. A. 109, 11372– 11377 (2012).

24. Tognoli, E. & Kelso, J. A. S. The Metastable Brain. Neuron 81, 35–48 (2014).

25. Freyer, F., Aquino, K., Robinson, P. A., Ritter, P. & Breakspear, M. Bistability and Non-Gaussian Fluctuations in Spontaneous Cortical Activity. J. Neurosci. 29, 8512–8524 (2009).

26. Freyer, F. et al. Biophysical Mechanisms of Multistability in Resting-State Cortical Rhythms. J. Neurosci. 31, 6353–6361 (2011).

27. Deco, G., Jirsa, V. K. & McIntosh, A. R. Emerging concepts for the dynamical organization of resting-state activity in the brain. Nat. Rev. Neurosci. 12, 43–56 (2011).

28. Nunez, P. L. & Cutillo, B. A. Neocortical Dynamics and Human EEG Rhythms. (Oxford University Press, 1995).

29. Greicius, M. D., Supekar, K., Menon, V. & Dougherty, R. F. Resting-State Functional Connectivity Reflects Structural Connectivity in the Default Mode Network. Cereb. Cortex 19, 72–78 (2009).

30. Roberts, J. A. et al. Metastable brain waves. Nat. Commun. 10, 1056 (2019).

31. Muller, L. et al. Rotating waves during human sleep spindles organize global patterns of activity that repeat precisely through the night. eLife 5, e17267 (2016).

32. Muller, L., Reynaud, A., Chavane, F. & Destexhe, A. The stimulus-evoked population response in visual cortex of awake monkey is a propagating wave. Nat. Commun. 5, 3675 (2014).

33. Robinson, P. A. et al. Eigenmodes of brain activity: Neural field theory predictions and comparison with experiment. NeuroImage 142, 79–98 (2016).

34. Gabay, N. C. & Robinson, P. A. Cortical geometry as a determinant of brain activity eigenmodes: Neural field analysis. *Phys*. Rev. E 96, 032413 (2017).

35. Henderson, J. A., Aquino, K. M. & Robinson, P. A. Empirical estimation of the eigenmodes of macroscale cortical dynamics: Reconciling neural field eigenmodes and resting-state networks. Neuroimage Rep. 2, 100103 (2022).

36. Gabay, N. C., Babaie-Janvier, T. & Robinson, P. A. Dynamics of cortical activity eigenmodes including standing, traveling, and rotating waves. *Phys*. Rev. E 98, 042413 (2018).

37. Tokariev, A. et al. Large-scale brain modes reorganize between infant sleep states and carry prognostic information for preterms. Nat. Commun. 10, 2619 (2019).

38. Chen, Y.-C. et al. The individuality of shape asymmetries of the human cerebral cortex. eLife 11, e75056 (2022).

39. Nunez, P. L. & Srinivasan, R. Electric Fields of the Brain: The Neurophysics of EEG. (Oxford University Press, 2006). doi:10.1093/acprof:oso/9780195050387.001.0001.

40. Bullmore, E. et al. Wavelets and functional magnetic resonance imaging of the human brain. NeuroImage 23 **Suppl 1**, S234–249 (2004).

41. Theiler, J., Eubank, S., Longtin, A., Galdrikian, B. & Farmer, J. D. Testing for nonlinearity in time series: the method of surrogate data. Phys. Nonlinear Phenom. 58, 77–94 (1992).

42. Koussis, N. & Group, S. N. SNG-Newy/eigenstrapping: v0.0.11 - Initial Release v3. Zenodo 10.5281/zenodo.10583040 (2024).

43. Pang, J. C. et al. Geometric constraints on human brain function. Nature 618, 566–574 (2023).

44. Hilbert, D. Methods of Mathematical Physics. (CUP Archive, 1985).

45. Robinson, P. A. et al. Determination of Dynamic Brain Connectivity via Spectral Analysis. Front. Hum. Neurosci. 15, (2021).

46. Blaser, R. & Fryzlewicz, P. Random Rotation Ensembles. J. Mach. Learn. Res. 17, 1–26 (2016).

47. Van Essen, D. C. et al. The WU-Minn Human Connectome Project: an overview. NeuroImage 80, 62–79 (2013).

48. Cohen, L. The generalization of the Wiener-Khinchin theorem. in *Proceedings of the 1998 IEEE International Conference on Acoustics, Speech and Signal Processing*, ICASSP’98 (Cat. No.98CH36181) vol. 3 1577–1580 vol.3 (1998).

49. Yura, H. T. & Hanson, S. G. Digital simulation of an arbitrary stationary stochastic process by spectral representation. JOSA A 28, 675–685 (2011).

50. Anselin, L. Local Indicators of Spatial Association—LISA. Geogr. Anal. 27, 93–115 (1995).

51. Vos de Wael, R., et al. BrainSpace: a toolbox for the analysis of macroscale gradients in neuroimaging and connectomics datasets. *Commun*. Biol. 3, 1–10 (2020).

52. Markello, R. D. et al. Standardizing workflows in imaging transcriptomics with the abagen toolbox. eLife 10, e72129 (2021).

53. Poldrack, R. et al. The Cognitive Atlas: Toward a Knowledge Foundation for Cognitive Neuroscience. *Front*. Neuroinformatics 5, (2011).

54. Yarkoni, T., Poldrack, R. A., Nichols, T. E., Van Essen, D. C. & Wager, T. D. Large-scale automated synthesis of human functional neuroimaging data. Nat. Methods 8, 665– 670 (2011).

55. Glasser, M. F. et al. The minimal preprocessing pipelines for the Human Connectome Project. NeuroImage 80, 105–124 (2013).

56. Margulies, D. S. et al. Situating the default-mode network along a principal gradient of macroscale cortical organization. Proc. Natl. Acad. Sci. 113, 12574–12579 (2016).

57. Allen, M. and P. Raincloud plots: a multi-platform tool for robust data visualization [version 2; peer review: 2 approved]. Wellcome Open Research vol. 4 (2021).

58. Miller, R. G. J. Simultaneous Statistical Inference. (Springer Science & Business Media, 2012).

59. Borne, L. et al. Functional re-organization of hippocampal-cortical gradients during naturalistic memory processes. NeuroImage 119996 (2023) doi:10.1016/j.neuroimage.2023.119996.

60. Buckner, R. L., Krienen, F. M., Castellanos, A., Diaz, J. C. & Yeo, B. T. T. The organization of the human cerebellum estimated by intrinsic functional connectivity. J. Neurophysiol. 106, 2322–2345 (2011).

61. Marquand, A. F., Haak, K. V. & Beckmann, C. F. Functional corticostriatal connection topographies predict goal-directed behaviour in humans. *Nat*. Hum. Behav. 1, 1–9 (2017).

62. Tian, Y., Margulies, D. S., Breakspear, M. & Zalesky, A. Topographic organization of the human subcortex unveiled with functional connectivity gradients. Nat. Neurosci. 23, 1421–1432 (2020).

63. Guell, X., Schmahmann, J. D., Gabrieli, J. D. & Ghosh, S. S. Functional gradients of the cerebellum. eLife 7, e36652 (2018).

64. Haak, K. V., Marquand, A. F. & Beckmann, C. F. Connectopic mapping with resting-state fMRI. NeuroImage 170, 83–94 (2018).

65. Campos Mantovanelli, B., Petry, M. T., Broetto Weiler, E. & Carlesso, R. Geostatistical interpolation based ternary diagrams for estimating water retention properties in soils in the Center-South regions of Brazil. Soil Tillage Res. 209, 104973 (2021).

66. Berman, T. D. & Allison, J. E. Coupling Thermomechanical Processing and Alloy Design to Improve Textures in Mg-Zn-Ca Sheet Alloys. JOM 73, 1450–1459 (2021).

67. Flores-Núñez, V. M. et al. Functional Signatures of the Epiphytic Prokaryotic Microbiome of Agaves and Cacti. Front. Microbiol. 10, (2020).

68. Maddess, T., Nagai, Y., James, A. C. & Ankiewcz, A. Binary and ternary textures containing higher-order spatial correlations. Vision Res. 44, 1093–1113 (2004).

69. Gupta, R., Mittal, A. & Patil, H. Robust order-based methods for feature description. 2010 IEEE Conference on Computer Vision and Pattern Recognition (CVPR) 334–341 (2010).

70. Koussis, N. & Group, S. N. SNG-Newy/eigenstrapping: v0.0.10 - Initial Release v2. Zenodo 10.5281/zenodo.10253593 (2023).

71. Breakspear, M., Brammer, M. & Robinson, P. A. Construction of multivariate surrogate sets from nonlinear data using the wavelet transform. Phys. Nonlinear Phenom. 182, 1– 22 (2003).

72. Breakspear, M., Brammer, M. J., Bullmore, E. T., Das, P. & Williams, L. M. Spatiotemporal wavelet resampling for functional neuroimaging data. Hum. Brain Mapp. 23, 1–25 (2004).

73. Breakspear, M. Dynamic models of large-scale brain activity. Nat. Neurosci. 20, 340–352 (2017).

74. Prichard, D. & Theiler, J. Generating surrogate data for time series with several simultaneously measured variables. Phys. Rev. Lett. 73, 951–954 (1994).

75. Aquino, K. M., Schira, M. M., Robinson, P. A., Drysdale, P. M. & Breakspear, M. Hemodynamic Traveling Waves in Human Visual Cortex. PLOS Comput. Biol. 8, e1002435 (2012).

76. Zalesky, A., Fornito, A., Cocchi, L., Gollo, L. L. & Breakspear, M. Time-resolved resting-state brain networks. Proc. Natl. Acad. Sci. 111, 10341–10346 (2014).

77. Choe, A. S. et al. Comparing test-retest reliability of dynamic functional connectivity methods. NeuroImage 158, 155–175 (2017).

78. Calhoun, V. D., Miller, R., Pearlson, G. & Adalı, T. The Chronnectome: Time-Varying Connectivity Networks as the Next Frontier in fMRI Data Discovery. Neuron 84, 262– 274 (2014).

79. Lurie, D. J. et al. Questions and controversies in the study of time-varying functional connectivity in resting fMRI. Netw. Neurosci. 4, 30–69 (2020).

80. Allen, E. A. et al. Tracking Whole-Brain Connectivity Dynamics in the Resting State. Cereb. Cortex 24, 663–676 (2014).

81. Menon, S. S. & Krishnamurthy, K. A Comparison of Static and Dynamic Functional Connectivities for Identifying Subjects and Biological Sex Using Intrinsic Individual Brain Connectivity. Sci. Rep. 9, 5729 (2019).

82. Lindquist, M. A., Xu, Y., Nebel, M. B. & Caffo, B. S. Evaluating dynamic bivariate correlations in resting-state fMRI: A comparison study and a new approach. NeuroImage 101, 531–546 (2014).

83. Aedo-Jury, F., Schwalm, M., Hamzehpour, L. & Stroh, A. Brain states govern the spatio-temporal dynamics of resting-state functional connectivity. eLife https://elifesciences.org/articles/53186/figures (2020) doi:10.7554/eLife.53186.

84. Nichols, T. E. & Holmes, A. P. Nonparametric permutation tests for functional neuroimaging: A primer with examples. Hum. Brain Mapp. 15, 1–25 (2002).

85. Bullmore, E. et al. Colored noise and computational inference in neurophysiological (fMRI) time series analysis: resampling methods in time and wavelet domains. Hum. Brain Mapp. 12, 61–78 (2001).

86. Patel, R. S., Van De Ville, D. & DuBois Bowman, F. Determining significant connectivity by 4D spatiotemporal wavelet packet resampling of functional neuroimaging data. NeuroImage 31, 1142–1155 (2006).

87. Wagner, H. H. & Dray, S. Generating spatially constrained null models for irregularly spaced data using Moran spectral randomization methods. Methods Ecol. Evol. 6, 1169– 1178 (2015).

88. Cao, T. et al. Mode-based morphometry: A multiscale approach to mapping human neuroanatomy. 2023.02.26.529328 Preprint at 10.1101/2023.02.26.529328 (2023).

89. Robinson, P. A., Rennie, C. J., Rowe, D. L., O’Connor, S. C. & Gordon, E. Multiscale brain modelling. Philos. Trans. R. Soc. B Biol. Sci. 360, 1043–1050 (2005).

90. Robinson, P. A., Rennie, C. J. & Rowe, D. L. Dynamics of large-scale brain activity in normal arousal states and epileptic seizures. Phys. Rev. E Stat. Nonlin. Soft Matter Phys. 65, 041924 (2002).

91. Breakspear, M. et al. A unifying explanation of primary generalized seizures through nonlinear brain modeling and bifurcation analysis. Cereb. Cortex N. Y. N 1991 16, 1296–1313 (2006).

92. Avena-Koenigsberger, A., Goñi, J., Solé, R. & Sporns, O. Network morphospace. J. R. Soc. Interface 12, 20140881 (2015).

93. Gollo, L. L. et al. Fragility and volatility of structural hubs in the human connectome. Nat. Neurosci. 21, 1107–1116 (2018).

94. Fischl, B., Sereno, M. I., Tootell, R. B. & Dale, A. M. High-resolution intersubject averaging and a coordinate system for the cortical surface. Hum. Brain Mapp. 8, 272–284 (1999).

95. Van Essen, D. C. et al. An integrated software suite for surface-based analyses of cerebral cortex. J. Am. Med. Inform. Assoc. JAMIA 8, 443–459 (2001).

96. Seo, S. & Chung, M. K. Laplace-Beltrami eigenfunction expansion of cortical manifolds. in 2011 IEEE International Symposium on Biomedical Imaging: From Nano to Macro 372–375 (2011). doi:10.1109/ISBI.2011.5872426.

97. Pang, J. C. et al. Reply to: Commentary on Pang et al. (2023) Nature. 2023.10.06.560797 Preprint at 10.1101/2023.10.06.560797 (2023).

98. Stewart, G. W. The Efficient Generation of Random Orthogonal Matrices with an Application to Condition Estimators. SIAM J. Numer. Anal. 17, 403–409 (1980).

99. Zito, T., Wilbert, N., Wiskott, L. & Berkes, P. Modular toolkit for Data Processing (MDP): a Python data processing framework. *Front*. Neuroinformatics 2, (2009).

100. Lehoucq, R. B., Sorensen, D. C. & Yang, C. ARPACK Users’ Guide: Solution of Large-Scale Eigenvalue Problems with Implicitly Restarted Arnoldi Methods. (SIAM, 1998).

101. Reuter, M., Wolter, F.-E., Shenton, M. & Niethammer, M. Laplace-Beltrami Eigenvalues and Topological Features of Eigenfunctions for Statistical Shape Analysis. Comput. Aided Des. 41, 739–755 (2009).

102. Wachinger, C. et al. BrainPrint: a discriminative characterization of brain morphology. NeuroImage 109, 232–248 (2015).

103. Vogel, J. W. et al. A molecular gradient along the longitudinal axis of the human hippocampus informs large-scale behavioral systems. Nat. Commun. 11, 960 (2020).

104. Royer, J. et al. Myeloarchitecture gradients in the human insula: Histological underpinnings and association to intrinsic functional connectivity. NeuroImage 216, 116859 (2020).

105. Glasser, M. F. et al. A multi-modal parcellation of human cerebral cortex. Nature 536, 171–178 (2016).

106. Makris, N. et al. Decreased volume of left and total anterior insular lobule in schizophrenia. Schizophr. Res. 83, 155–171 (2006).

107. Frazier, J. A. et al. Structural brain magnetic resonance imaging of limbic and thalamic volumes in pediatric bipolar disorder. Am. J. Psychiatry 162, 1256–1265 (2005).

108. Desikan, R. S. et al. An automated labeling system for subdividing the human cerebral cortex on MRI scans into gyral based regions of interest. NeuroImage 31, 968– 980 (2006).

109. Goldstein, J. M. et al. Hypothalamic abnormalities in schizophrenia: sex effects and genetic vulnerability. Biol. Psychiatry 61, 935–945 (2007).

110. Karras, T., et al. Analyzing and Improving the Image Quality of StyleGAN. Preprint at 10.48550/arXiv.1912.04958 (2020).

111. Tan, X. & Triggs, B. Enhanced Local Texture Feature Sets for Face Recognition Under Difficult Lighting Conditions. in Analysis and Modeling of Faces and Gestures (eds. Zhou, S. K., Zhao, W., Tang, X. & Gong, S.) 168–182 (Springer, Berlin, Heidelberg, 2007). doi:10.1007/978-3-540-75690-3_13.

