## Supplementary Materials for "Generation of surrogate brain maps preserving spatial autocorrelation through random rotation of geometric eigenmodes"

### S1. Treatment of the residuals of the eigenmode decomposition

The eigenmode decomposition,

$$\begin{aligned} y(\boldsymbol{x})=\sum_{\Lambda=0}^{G} \sum_{\mu=-\Lambda}^{\Lambda} \left( \beta_{\Lambda\mu}\psi_{\Lambda\mu}\left( \boldsymbol{x} \right) \right)+\epsilon_{G}\left( \boldsymbol{x} \right),\#\left( S1 \right) \end{aligned}$$

on a mesh with *N* vertices on discrete surface $\boldsymbol{x}$ is complete when a total of *N* eigenmodes are used, whereby the residual term ε limits to zero (to numerical accuracy). However, for a highly resolved cortical mesh (e.g., 32,492 vertex points per hemisphere), a complete representation of *n=*32,492 modes per hemisphere carries a substantial computational burden. As power diminishes at higher spatial frequencies, incomplete decompositions typically yield unstructured residual error for *n*<<*N*. Residuals are calculated by subtracting the reconstructed data from the original data. Despite their relatively small coefficients, they contain substantial variance that needs to be included in the surrogate data (Fig. S2C). These residuals can be added back into the surrogates directly, resulting in better matching of the variogram even when resampling a small number of modes (*n* ~ 200; Fig. S2A). However, this re-addition of the original residual leads to sub-optimal behavior of the resulting surrogate maps, such that they are correlated with the original data and each other (Fig. S2B), due to the perfect matching of these small terms between the original and all the surrogate maps.

To retain the variance without yielding pairwise correlated surrogates, the residuals can be *permuted* before being added to the surrogates (Fig. S3). However, the combined effects of adding the permuted residuals and undertaking amplitude adjustment (see below) only preserves SA if the number of modes is sufficiently high (Fig. S3D).

### S2. Amplitude adjustment

Amplitude adjustment (AA) is commonly used in Fourier decomposition and wavelet-based surrogate techniques for time-series analysis when the original data does not possess a normal amplitude distribution ^1–4^. This step is required to preserve the original distribution of the data, because, due to the Central Limit Theorem, resampling methods yield surrogate data with a Gaussian amplitude distribution, hence causing a mismatch with empirical data sets that have non Gaussian amplitude distribution ^1,2,5,6^ (Fig. 1, panel C). AA is performed by rank-sorting the surrogate data and the empirical data, then replacing the highest-ranked values of the surrogate data with the highest-ranked values of the empirical data, then the next highest, and so on, until the surrogate map has the same amplitude distribution as the original.

As evident in Fig. S3, the AA has a small but non-negligible impact on the variogram. This subtle whitening effect (increasing the variance at small separations) is a known effect when AA is applied to Fourier or wavelet-based resampling techniques and arises due to the re-sorting and re-insertion of the original data samples ^1–3,7^. In the present setting, this effect means that the final number of modes, the application of AA and the need to permute the residuals before re-insertion are somewhat co-dependent. These dependencies can be mitigated when using a sufficient number of modes to generate surrogate data. In our testing we recommend a conservative estimate of minimum 200 modes for standard MRI datasets with voxel resolutions > 1mm and smoothing kernels of 6 mm full-width half-maximum and greater.

### S3. HCP data

All preprocessed fMRI data was accessed from the Human Connectome Project ^8^. No further preprocessing steps were applied, and data was analyzed from 255 unrelated healthy individuals (aged 22-35 years, 132 females and 123 males). This is the largest cohort of the HCP excluding twins or siblings that had completed all tasks and resting-state acquisitions. All procedures entailed in this study were carried out in accordance with local ethics guidelines and with approval from the local ethics committee (University of Newcastle HREC ref: H-2020-0443). Image acquisition parameters, task protocols, and preprocessing pipelines are thoroughly detailed in refs. ^8,9^.

Seven task domains detailed in Supplementary Table 2 and task-free resting-state fMRI were analyzed in our study, already preprocessed by HCP. Volumetric contrast activation maps for tasks were resampled to *fs-LR-32k* CIFTI space using Connectome Workbench tools. Volumetric timeseries for resting-state data were used for the cortico-subcortical connectivity gradients. Further information on the construction of these gradients can be found in Supplementary Information-S7.

### S4. Gaussian random fields

GRF pairs were generated by adapting Python code from <https://github.com/markello_spatialnulls/parspin>, using the method described in ^10^. First, two uniformly spaced three-dimensional grids with tiling corresponding to the dimensions of the MNI152 2mm standard volume (91x107x91) were generated. At each point on the grid pairs, a random sample was drawn from a multivariate normal distribution tuned to correlate across grids at |*r*| = 0.15±0.005. This shortens the time to randomly generate pairs that correlate when SA is low ($\alpha$ $\cong$ 0). This is because GRF pairs with SA have a high chance of being correlated, while the random chance of white noise GRF pairs being correlated is very low ^11^.

Each pair was then normalized to have zero mean and unit variance. The normalized pairs were then projected to the *fsaverage5* surface with 10,242 vertices by FreeSurfer *mri_vol2surf*. The correlations of the resulting pairs were then checked after the medial wall was removed and discarded if |*r*| did not fall between 0.145 and 0.155.

The histogram of correlations between random pairs of GRFs for each $\alpha$ from 0.0 to 4.0 are provided in Fig. S4. This shows that while the correlations are zero-centered, the tails of the distributions widen as $\alpha$ increases.

### S5. Moran’s *I*

Moran’s *I* is a measure of the autocorrelation of spatial data, commonly utilized in geostatistics and texture analysis ^12,13^. Unlike the variogram, Moran’s *I* is not distance dependent, but rather a composite of SA across all pairwise distances.

Moran’s *I* of a function $f_{i}=f(\boldsymbol{x}_{\boldsymbol{i}})$ on a discrete surface $\boldsymbol{x}_{\boldsymbol{i}}$ with distance weights $w_{ij}=1/\left| \boldsymbol{x}_{\boldsymbol{i}}-\boldsymbol{x}_{\boldsymbol{j}} \right|$is defined as,

$$\begin{aligned} I=\frac{N}{W}\frac{\sum_{i=1}^{N} \sum_{j=1}^{N} w_{ij}\left( f_{i}-\bar{f} \right)(f_{j}-\bar{f})}{\sum_{i=1}^{N} \left( f_{i}-\bar{f} \right)^{2}},\#\left( S2 \right) \end{aligned}$$

where $N$ is the total number of vertices on ***x***, $\bar{f}$ is the mean of $f$, $w_{ij}$ are the elements of distance weights where $w_{i=j,j=i}=0$ and $W=\sum_{i=1}^{N} \sum_{j=1}^{N} w_{ij}$ ^13^.

We benchmarked the SA-replicating property of eigenstrapping (blue) against BrainSMASH (green) and Spin Test (yellow) in Fig. S5. Evaluating the performance of each method was achieved by calculating the change in the Moran’s $I \left( \Delta I=empirical I-null I \right)$ statistic between 1000 simulated maps (GRFs) and 1000 surrogates for each value of $\alpha$ across a range from 0.0 – 4.0 (Fig. S5). A pairwise distance matrix with shape (10242x10242) was derived for the surfa–e that all GRFs were mapped to (*fsaverage5*; see Supplementary Information-S5) and inverted to derive distance weights (the diagonal was set to zero). This weight matrix was then used to calculate Moran’s *I*.

Low values of $\alpha$ (0.0 and 1.0) yielded spatial maps with little SA and hence the Moran’s *I* surrounded zero (cyan; Fig. S5A) or was slightly positive (Fig. S5B). All three surrogate methods performed reasonably in this range, with the change in Moran’s *I* ($\Delta I$) centered at zero. Higher values of $\alpha$ generated smoother spatial maps with a corresponding increase in Moran’s I (Fig. S5C-E). Notably, surrogates derived using the BrainSMASH method are considerably whiter than the original maps (Moran’s *I* lower than the source map, green). Both eigenstrapping and the spin test preserve the smoothness appropriately – the difference in Moran’s *I* is centered around zero (blue and yellow). Raw Moran’s *I* values are plotted accordingly in Fig. S6 to show the notable decrease in SA of BrainSMASH surrogates.

### S6. Calculation of cortico-subcortical functional connectivity gradients

Connection topographical (*connectopic*) patterns (known as gradients) of resting-state activity capture the similarity of the functional connectivity patterns of neighbouring voxels and possess complex SA^14^. Calculation of these cortico-subcortical functional connectivity gradients followed a previously published methodology ^14^. Cortico-subcortical functional connectivity gradients were derived by correlating the timeseries of BOLD in the subcortical mask (either thalamus, hippocampus, or striatum) with the principal components (PCs) in whole brain gray matter (GM). Subcortical masks were derived by binarizing the Harvard-Oxford subcortical atlas at 25% probability for each region. PCs were calculated by singular-value decomposition (SVD) of the timeseries, of shape ($T\times T-1$, where $T$ is the number of volumes). PCs were then correlated with the columns of subcortical activity over time (of shape $(N\times T$) where $N$ is the number of voxels in subcortical GM). The correlation of cortical SVD components and subcortical voxels results in a correlation matrix $\boldsymbol{C}$ of shape $\left( N\times T-1 \right)$. This correlation matrix was then Fisher’s Z-transformed, yielding a positive normal distribution of values between 0 and 1. We characterized the functional connectivity similarity of every subcortical voxel to every other subcortical voxel by deriving the $\eta^{2}$ coefficient row-wise of $\boldsymbol{C}$, resulting in a symmetric similarity matrix $\boldsymbol{S}$,

$$\begin{aligned} S_{\alpha,\beta}=1-\frac{\sum_{j=1}^{p} \left[ \left( C_{\alpha,j}-\mu_{j} \right)^{2}-\left( C_{\beta,j}-\mu_{j} \right)^{2} \right]}{\sum_{j=1}^{p} \left[ \left( C_{\alpha,j}-\bar{\mu}_{j} \right)^{2}-\left( C_{\beta,j}-\bar{\mu}_{j} \right)^{2} \right]},\#\left( S3 \right) \end{aligned}$$

where the matrix $\boldsymbol{S}$ has rows and columns $\left( \alpha,\beta\right)$ of shape $\left( V\times V \right)$. $V$ is the number of voxels in the subcortex, $j$ is the column of the correlation matrix $\boldsymbol{C}$, $p$ corresponds to the total number of columns (the SVD-components), $\mu_{j}=\frac{C_{\alpha,j}-C_{\beta,j}}{2}$, and $\bar{\mu}$ is the mean of all $\mu$ across all $p$ SVD-components.

To derive connectopic gradients, the similarity matrix $\boldsymbol{S}$ must first be made into sparse matrix $\boldsymbol{W}$ to calculate the graph Laplacian. $\boldsymbol{S}$ was rendered sparse according to the following rule for each element of $\boldsymbol{W}$,

$$\begin{aligned} W_{i,j}=\left\{ \begin{matrix} S_{i,j} \text{if}\left| \left| S_{i}-S_{j} \right| \right|^{2} <\varepsilon\\ 0 \text{if}\left| \left| S_{i}-S_{j} \right| \right|^{2} \geq\varepsilon\end{matrix} \right.,\#\left( S4 \right) \end{aligned}$$

where $\varepsilon$ is the minimum value required for the graph to remain connected. The graph Laplacian $\boldsymbol{L}$ is then calculated by $\boldsymbol{L}=\boldsymbol{D}-\boldsymbol{W},$ where $\boldsymbol{D}$ is equal to the trace of $\boldsymbol{W}$**.** The eigenvalues and eigenvectors were derived from the generalized eigenvalue problem,

$$\begin{aligned} \boldsymbol{Lu}=-\zeta\boldsymbol{Du},\#\left( S5 \right) \end{aligned}$$

where $\Delta$ is the Laplacian operator, $\zeta$ are the eigenvalues, and $\boldsymbol{u}$ are the connectopic eigenfunctions corresponding to the eigenvalues. These eigenfunctions are maps wherein voxels with similar values have similar connectivity patterns, and voxels with different values have different connectivity patterns ^14–16^. The SA of these patterns reflects the typically gradual change across cortex and subcortex of these functional connectivity patterns, with abrupt changes limited to functional boundaries ^16^.

### S7. Treatment of the medial wall in the Spin Test

Rotation of the cortical surface on the sphere results in a substantial number of “medial wall” vertices labeled as non-data (or *unknown*, as FreeSurfer labels it) being re-located to the cortex, with the same number of cortical vertices being rotated onto the medial wall.

There are three general solutions to this problem: (1) Masking out any vertices that are lost (*i.e*., old medial wall plus new medial wall), that is removing them from any proceeding analyses. This can remove up to 20-30% of the total number of samples on the surface. (2) Setting all values that lie within the medial wall to zero. (3) Interpolating values across medial wall vertices that are rotated into the new cortical surface. All three solutions bias further analyses to varying degrees, though the most widely used option is (1), which is what this study used to compare in the main paper. The presence of missing data is indicated by the black marker in Fig. 4A.

Eigenstrapping surmounts this issue by setting a boundary around the medial wall and limiting the LBO operator to the ensuing closed surface across the cortex. This ensures that rotations are performed in the modal space $\boldsymbol{z}$, not vertex space.

### S8. Surrogate computation time

The time taken for one CPU/thread to compute 1000 surrogates on several different levels of surface density (from volumetric and surface-based analyses), with and without pre-computed eigenmodes is provided in Fig. S9 (specifications for the computational device are detailed in Supplementary Table S3).

For subcortical maps (*volumetric*), precomputing and caching eigenmodes makes no difference in computation time (“no caching” and “with caching”). In the two cortical surfaces used in this study (*fs-LR-32k* and *fsaverage5*, with 32,492 and 10,242 vertices respectively), the computation of eigenmodes slightly increases the computation time, particularly on the denser *fs-LR-32k* surface, adding on average 50 seconds to the runtime. When the modes have been pre-computed, the two surfaces take on average 250 seconds and 1200 seconds to compute 1000 surrogates. Deriving more modes requires more computation time as the complexity of the mode calculation increases linearly with the surface size ^17^.

The most computationally intensive component of the algorithm (Alg. 1) is a *for* loop that generates $n\times n$ random rotation matrices and performs dot product operations on these. Here, the complexity is no greater than $\mathcal{O}\left( n^{2} \right)$, where $n$ is the number of modes in the largest group (the size of the largest group in the first 10,000 modes is 201; Fig. S9). This simplifies even further to $\mathcal{O}\left( N \right)$ (where $N$ is the number of vertices) when the rotated modes have been pre-computed. Future development of the method will seek to provide openly available sources of rotated modes both for speeding up computation and reproducibility.

To overcome the challenge of computation of modes for standard surfaces, a selection of pre-computed modes are integrated into the open-release Python package. The precomputation of modes for standard surfaces should be suitable for the end-user in the majority of cases, as group-level neuroimaging analyses (where null hypothesis testing would generally occur) require registration to a standard space.

### Supplementary References

1.	Schreiber, T. & Schmitz, A. Improved Surrogate Data for Nonlinearity Tests. *Phys. Rev. Lett.* **77**, 635–638 (1996).

2.	Schreiber, T. & Schmitz, A. Surrogate time series. *Phys. Nonlinear Phenom.* **142**, 346–382 (2000).

3.	Lancaster, G., Iatsenko, D., Pidde, A., Ticcinelli, V. & Stefanovska, A. Surrogate data for hypothesis testing of physical systems. *Phys. Rep.* **748**, 1–60 (2018).

4.	Breakspear, M., Brammer, M. & Robinson, P. A. Construction of multivariate surrogate sets from nonlinear data using the wavelet transform. *Phys. Nonlinear Phenom.* **182**, 1–22 (2003).

5.	Breakspear, M., Brammer, M. J., Bullmore, E. T., Das, P. & Williams, L. M. Spatiotemporal wavelet resampling for functional neuroimaging data. *Hum. Brain Mapp.* **23**, 1–25 (2004).

6.	Patel, R. S., Van De Ville, D. & DuBois Bowman, F. Determining significant connectivity by 4D spatiotemporal wavelet packet resampling of functional neuroimaging data. *NeuroImage* **31**, 1142–1155 (2006).

7.	Breakspear, M., Brammer, M. J., Bullmore, E. T., Das, P. & Williams, L. M. Spatiotemporal wavelet resampling for functional neuroimaging data. *Hum. Brain Mapp.* **23**, 1–25 (2004).

8.	Van Essen, D. C. *et al.* The WU-Minn Human Connectome Project: an overview. *NeuroImage* **80**, 62–79 (2013).

9.	Glasser, M. F. *et al.* The minimal preprocessing pipelines for the Human Connectome Project. *NeuroImage* **80**, 105–124 (2013).

10.	Markello, R. D. & Misic, B. Comparing spatial null models for brain maps. *NeuroImage* **236**, 118052 (2021).

11.	Burt, J. B., Helmer, M., Shinn, M., Anticevic, A. & Murray, J. D. Generative modeling of brain maps with spatial autocorrelation. *NeuroImage* **220**, 117038 (2020).

12.	Moran, P. A. P. Notes on Continuous Stochastic Phenomena. *Biometrika* **37**, 17–23 (1950).

13.	Anselin, L. Local Indicators of Spatial Association—LISA. *Geogr. Anal.* **27**, 93–115 (1995).

14.	Haak, K. V., Marquand, A. F. & Beckmann, C. F. Connectopic mapping with resting-state fMRI. *NeuroImage* **170**, 83–94 (2018).

15.	Marquand, A. F., Haak, K. V. & Beckmann, C. F. Functional corticostriatal connection topographies predict goal-directed behaviour in humans. *Nat. Hum. Behav.* **1**, 1–9 (2017).

16.	Tian, Y., Margulies, D. S., Breakspear, M. & Zalesky, A. Topographic organization of the human subcortex unveiled with functional connectivity gradients. *Nat. Neurosci.* **23**, 1421–1432 (2020).

17.	Lehoucq, R. B., Sorensen, D. C. & Yang, C. *ARPACK Users’ Guide: Solution of Large-Scale Eigenvalue Problems with Implicitly Restarted Arnoldi Methods*. (SIAM, 1998).

18.	Robinson, P. A. *et al.* Eigenmodes of brain activity: Neural field theory predictions and comparison with experiment. *NeuroImage* **142**, 79–98 (2016).

19.	Pang, J. C. *et al.* Reply to: Commentary on Pang et al. (2023) Nature. 2023.10.06.560797 Preprint at https://doi.org/10.1101/2023.10.06.560797 (2023).

### Supplementary Figures

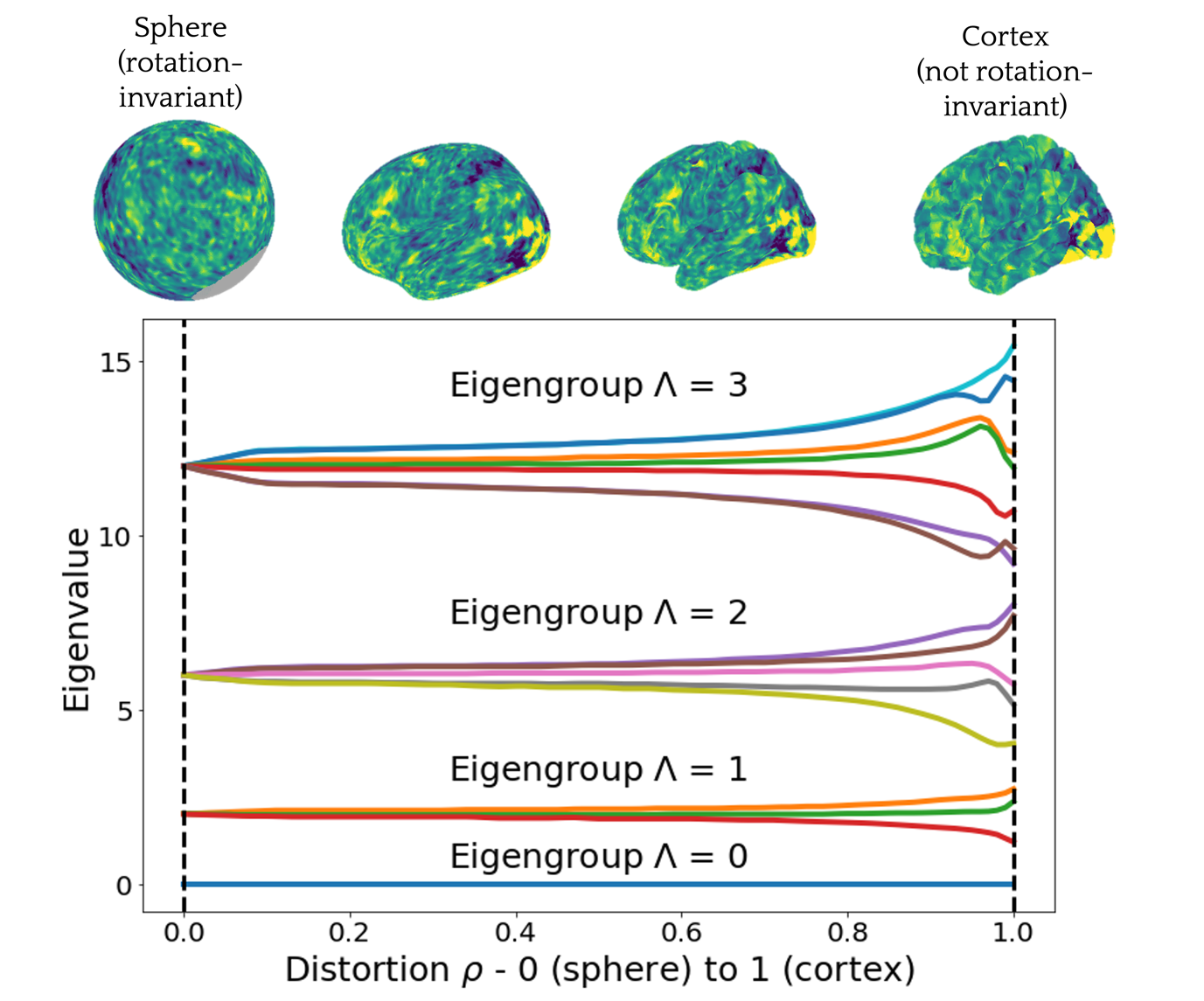

**Fig. S1.** Eigenvalue spectrum of the first 4 eigengroups on the left hemisphere, corresponding to the first 16 eigenvalues. The spectrum is evaluated at 100 points as the cortical surface ($\rho$ = 1.0) is gradually inflated to the sphere ($\rho$ = 0.0). The degree of distortion is proportional to the folding of the cortex. Note that the inflation of the surface was fixed so that surface area was kept consistent across surfaces. Eigenvalue figure adapted with permission from ^18,19^.

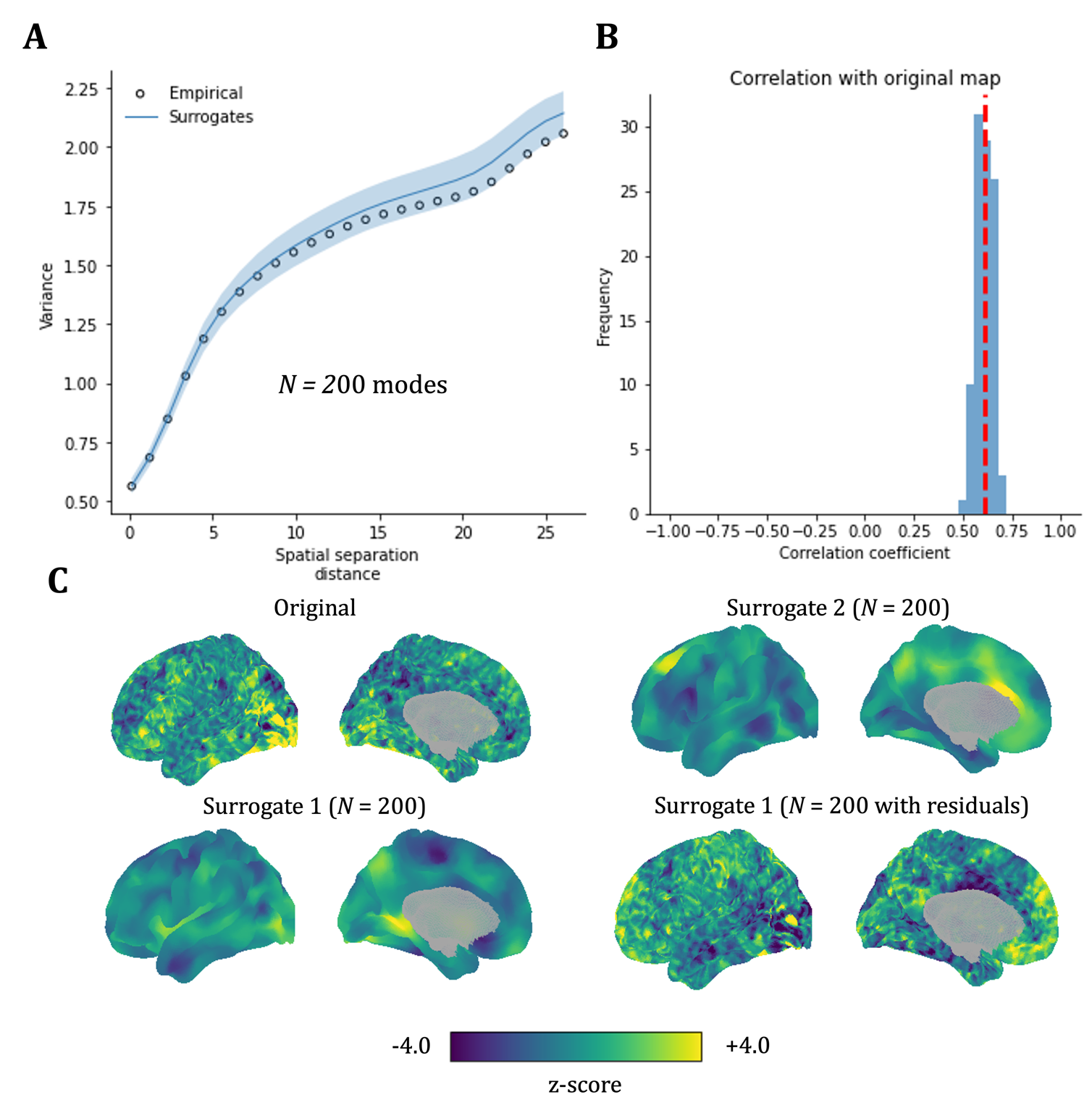

**Fig. S2.** The effect of including model residuals in surrogate brain maps. (A) Variogram of HCP task contrast data with N = 200 modes for modal decomposition. Note the closely matching variograms of the 1000 surrogates with original residuals added back to data after Eigenstrapping. (B) Including the original residuals in the surrogate distribution induces a strong correlation of the surrogates with each other and with the original data. Red dashed line is the average correlation of the surrogates with the empirical data. (C) Original map plotted alongside exemplar surrogates at N = 200. Notice the smoothness of the surrogates with low modes – these are then “recolored” by addition of the residuals to match the original, but this shifts the resulting distribution to highly correlated values unless the residuals are randomly permuted prior to inclusion.

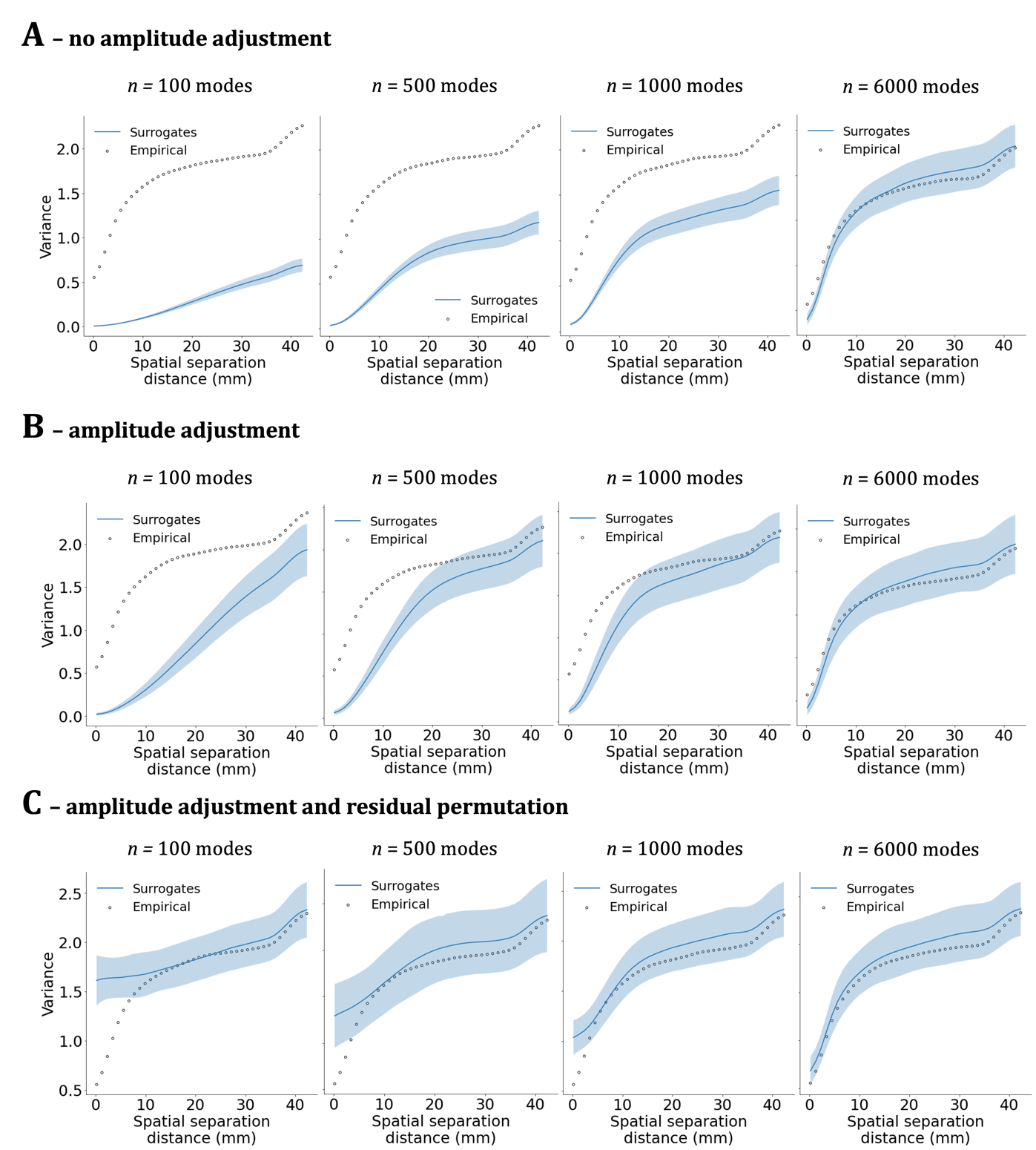

**Fig. S3.** Variograms of HCP *emotion* task data (see Supplementary Table 2) show the fit in (A) without amplitude adjustment is lower than in (B) where amplitude adjustment improves the fit of the surrogates (blue line) via a slight whitening effect, as well as broadening the standard deviation of the variograms (blue shaded area). Permuting the residuals increases the (zero-lag) variance of the surrogates in (C), which decreases to be in line with the empirical zero-lag as the number of modes increase.

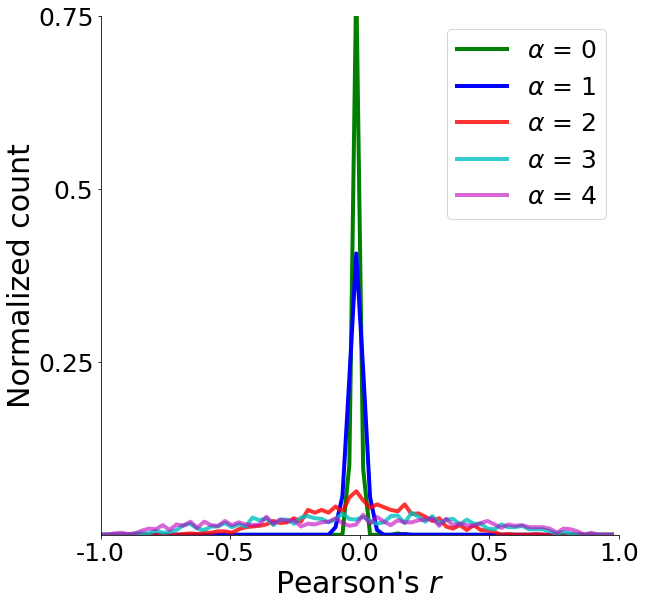

**Fig. S4.** Distribution of Pearson correlation between randomly paired generated maps as a function of $\alpha$. Distributions are shown for $\alpha$=0 (green), $\alpha$=1 (blue), $\alpha$=2 (orange), $\alpha$=3 (cyan), and $\alpha$=4 (magenta). Note that values are normalized by the total number of pairs for each $\alpha$ (1000).

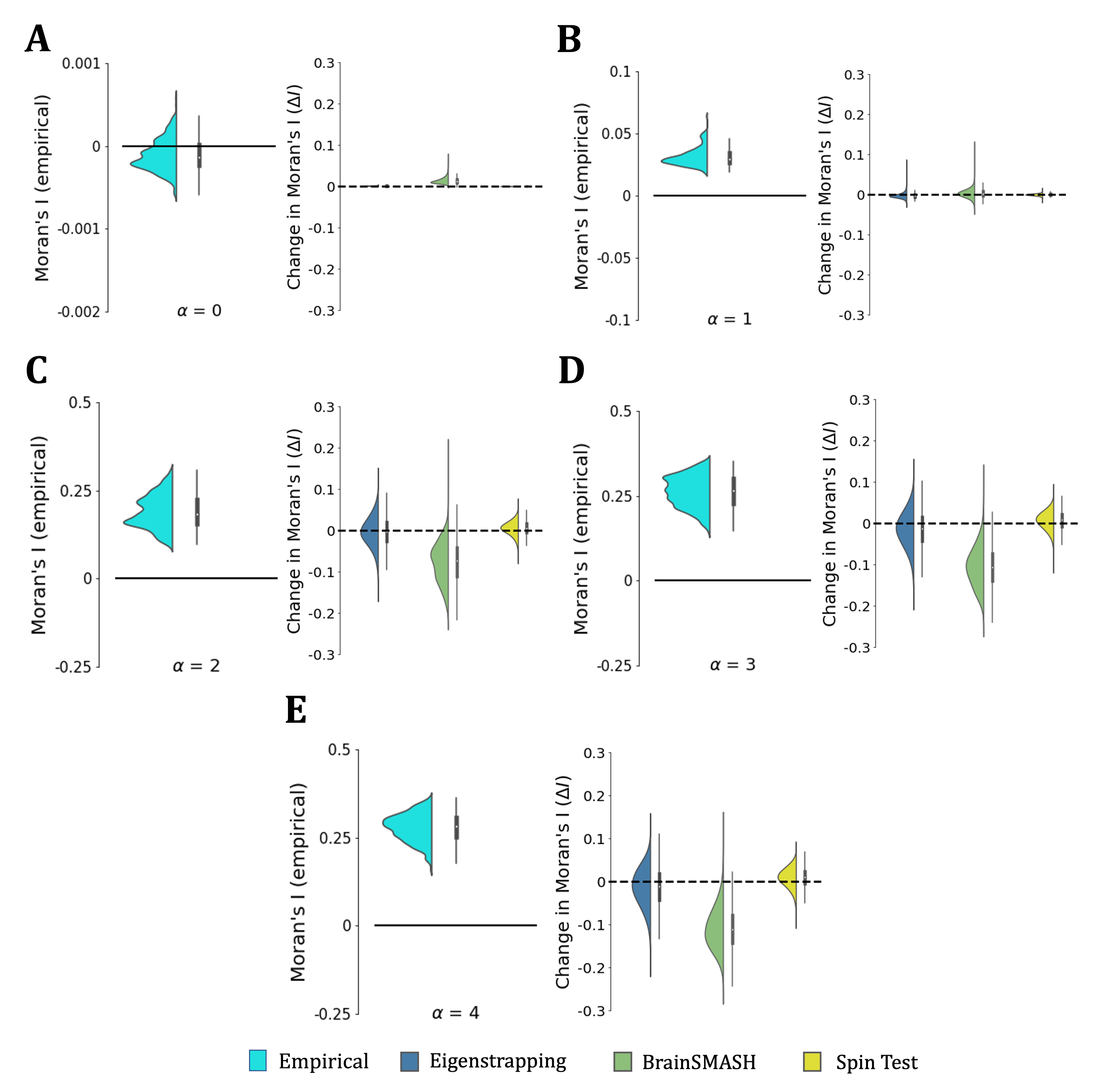

**Fig. S5.** Moran’s $I$ statistic computed on 1000 GRFs (left in light green) as a function of increasing SA. A: $\alpha$ = 0.0; B: $\alpha$ = 1.0; C: $\alpha$ = 2.0; D: $\alpha$ = 3.0; E: $\alpha$ = 4.0. Change in Moran’s $I$ statistic ($\Delta$I; empirical$-$null) is shown with rainclouds for eigenstrapping in blue, BrainSMASH in green, and Spin Test in yellow. Raw Moran’s I values for empirical and surrogate classes for each $\alpha$ are given in Supplementary Fig. S6.

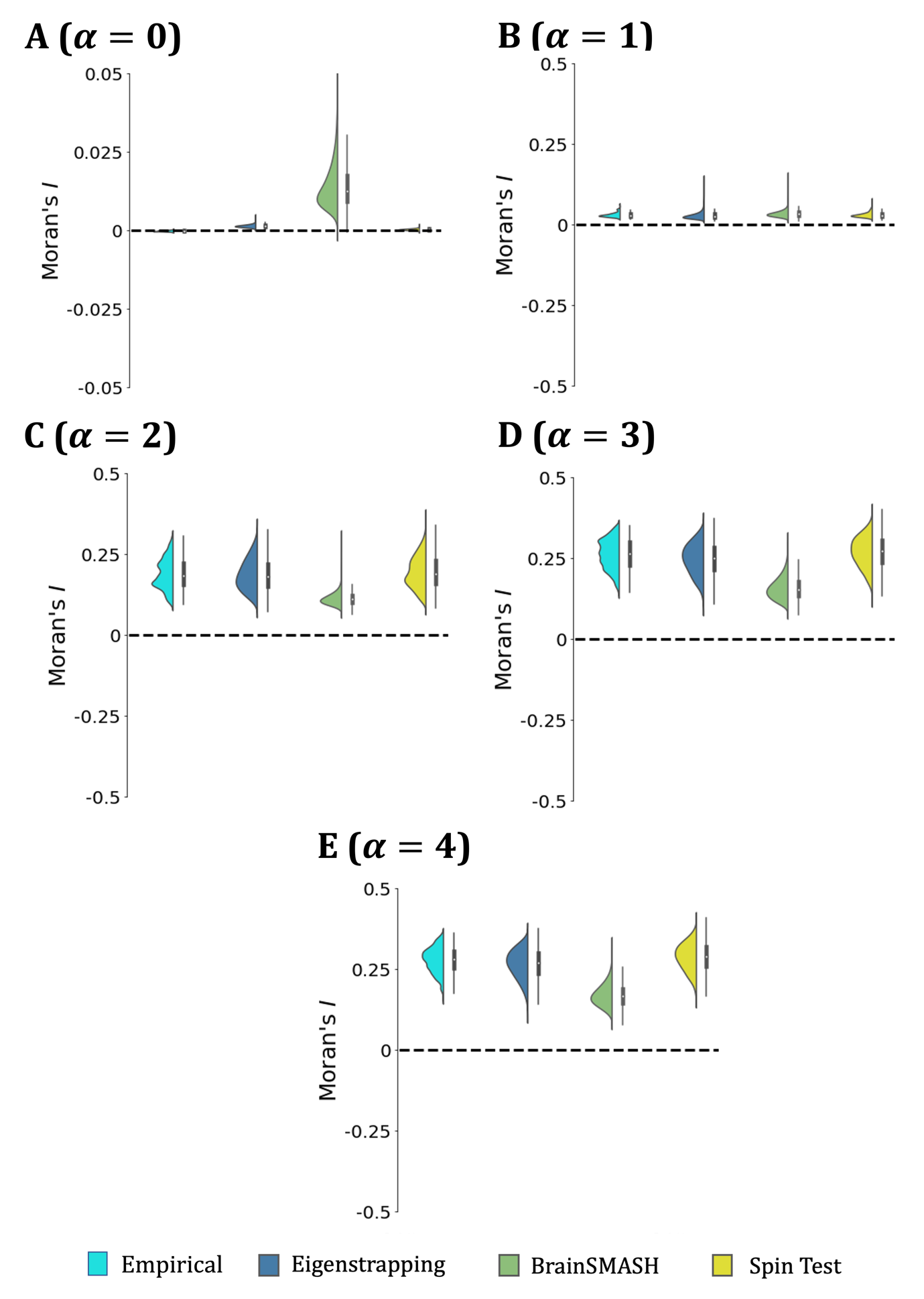

**Fig. S6.** Moran’s *I* as a function of spatial autocorrelation $\alpha$: (A) $\alpha=0$, (B) $\alpha=1$, (C) $\alpha=2$, (D) $\alpha=3$, (E) $\alpha=4$. Empirical values from simulated GRFs (1000 in each $\alpha$) are plotted in cyan histograms; Moran’s *I* of 1000 eigenstrapping surrogates of each GRF in blue; Moran’s *I* of 1000 BrainSMASH surrogates of each GRF in green; Moran’s *I* of 1000 Spin Test surrogates of each GRF in yellow.

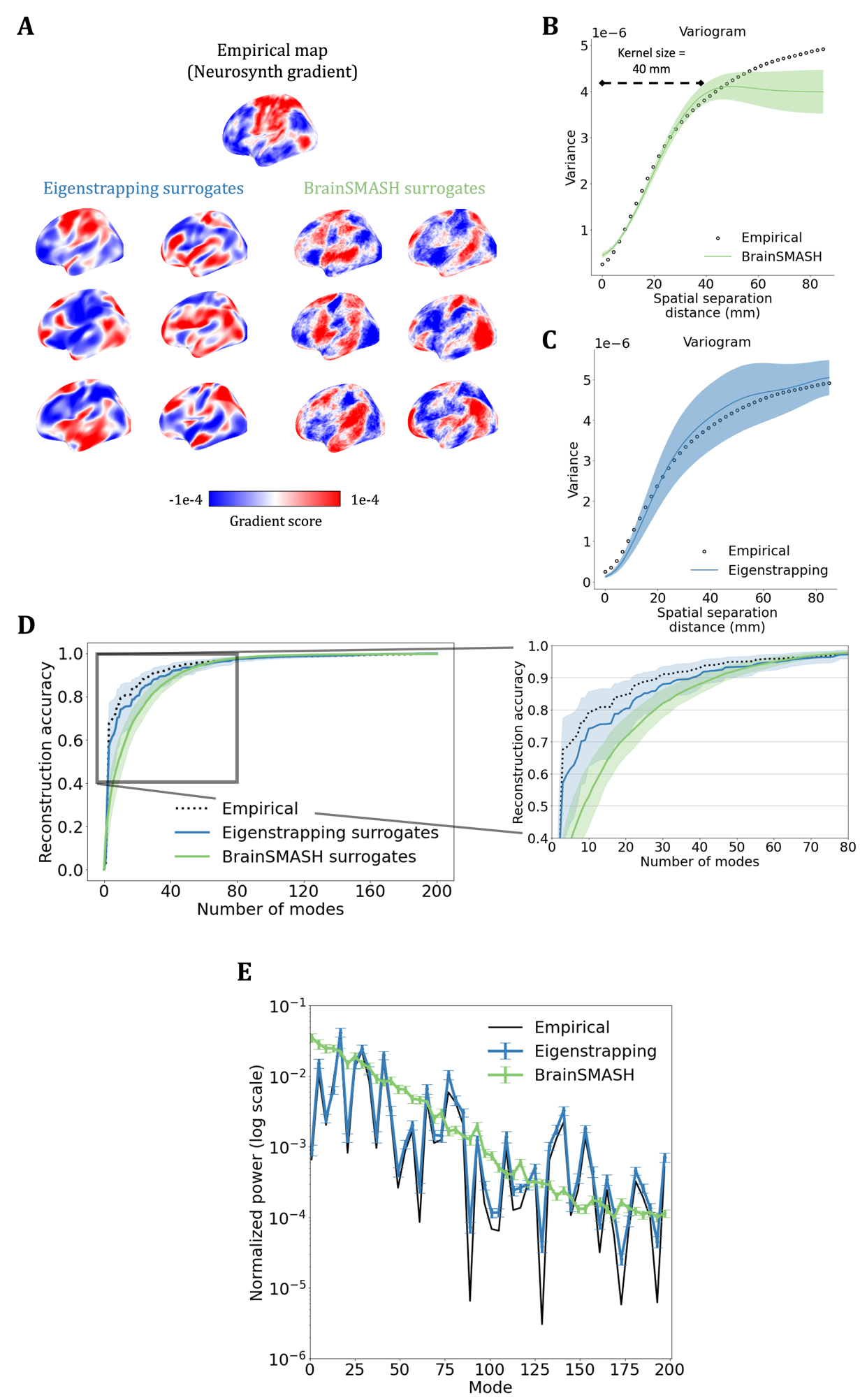

**Fig. S7.** Impact of BrainSMASH on spatial autocorrelation. (A) Empirical cognitive brain map (NeuroSynth cognitive terms 1st principal gradient) from Fig. 4 is shown above six exemplar surrogates in each method. The whitening of the BrainSMASH surrogates is evident as the increased speckling. (B) The whitening effect is strongest at separations wider than BrainSMASH kernel (40 mm kernel). There also exists a slight whitening at the zero-lag variance (the intercept of the variogram). (C) Eigenstrapping replicates the variogram to very long ranges (>80 mm), with a greater variance than BrainSMASH at 40-60 mm. The mean variance of the surrogates follows the empirical curve across the entire spectrum. (D) Reconstruction of the empirical data (black dashed line), the eigenstrapped surrogates (blue), and BrainSMASH surrogates (green). The eigenstrapping surrogates show similar reconstruction to the original data. The inset panel shows the slower reconstruction accuracy of the BrainSMASH surrogates. (E) The average power spectrum of eigenstrapping surrogates (blue) is nearly identical to the empirical power spectrum (Pearson’s *r* = 0.961). The average power spectrum of BrainSMASH surrogates (green) reproduces the slope, but not the variability of the empirical spectrum (Pearson’s *r* = 0.401). Error bars in panel E denote standard error.

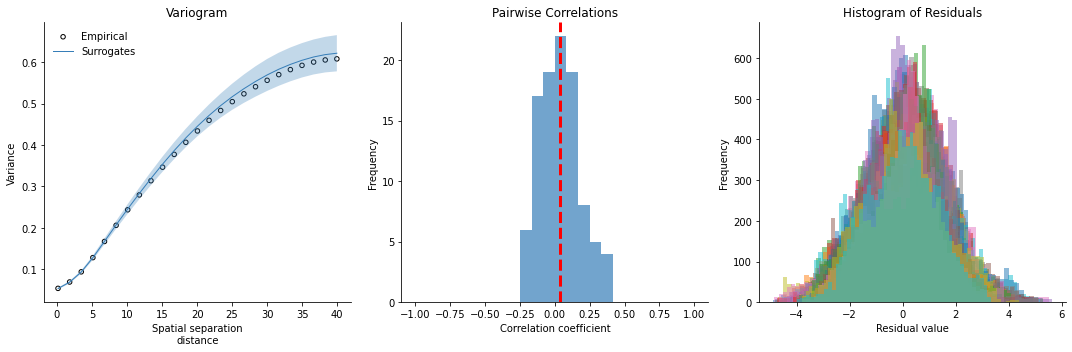

**Fig. S8.** Eigenstrapping diagnostic tools for the end-user. Left panel: Variogram of original data against surrogates. Middle panel: Pairwise correlations of surrogates with original data. Right panel: Histogram of residuals calculated from subtracting surrogates from original data.

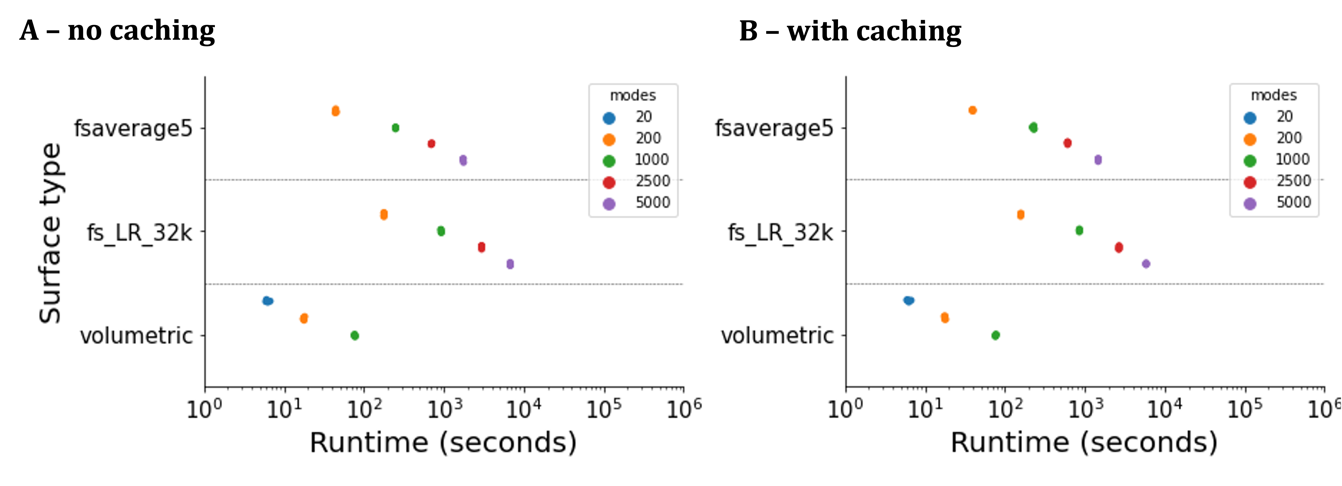

**Fig. S9.** Computation runtime for 1,000 surrogates. The computation time for the *fsaverage* (blue), *fs-LR-32k* (orange), and *volumetric* (green) resolution data. Each null was run with different numbers of modes (20: blue; 200: orange; 1,000: green; 2,500: red; purple: 5,000), five times on a simulated brain map ($\alpha=2.0$) for *fsaverage*; HCP task contrast data for *fs-LR-32k*; HCP cortico-subcortical gradient in the striatum for *volumetric*, consisting of 2,230 vertices). Repeats are plotted as separate dots. (A) The computation time of 1,000 surrogates with no pre-computation of modes. (B) The computation time of surrogates with pre-computation of modes. All computations were performed using one CPU/thread. Specifications for the computational device are listed in Supplementary Table 3.

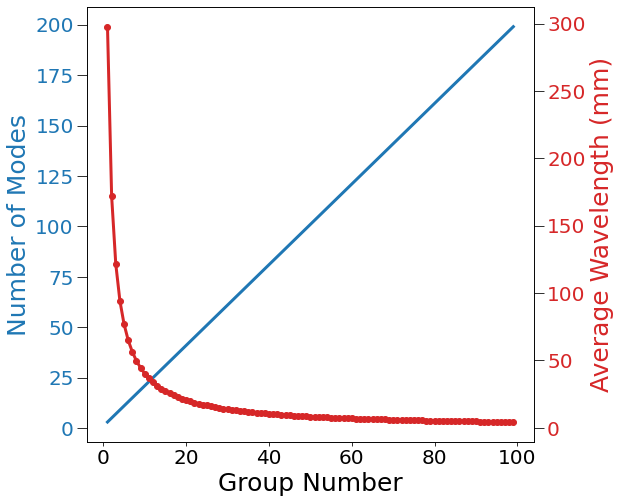

**Fig. S10.** Number of eigenmodes per eigengroup (blue) and wavelength in mm of each eigengroup (red) up to the first 100 eigengroups (corresponding to the first 10000 modes).

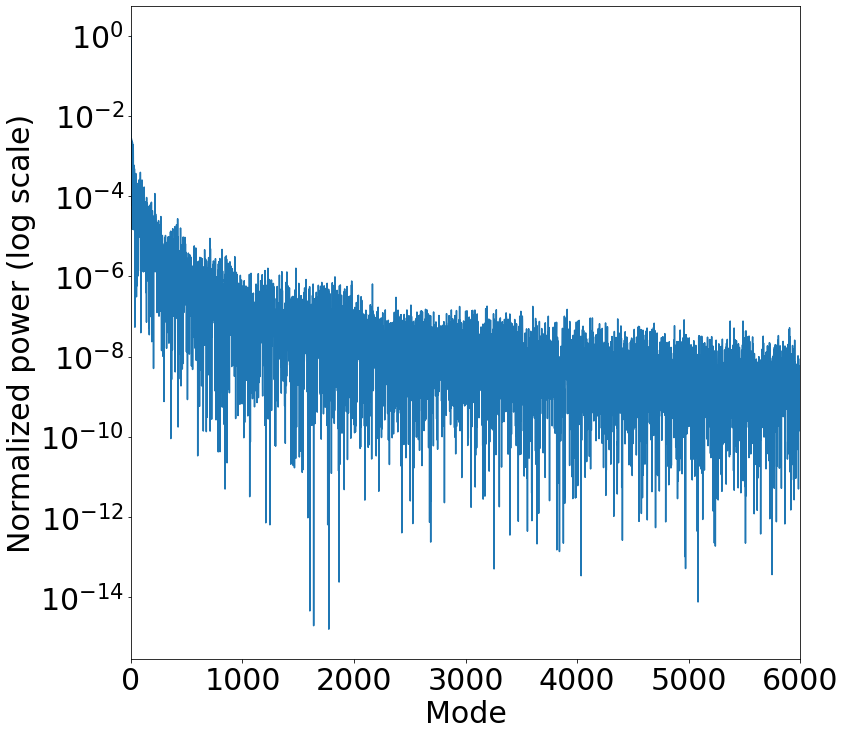

**Fig. S11.** Normalized eigenmode power spectral density derived from 6000 mode coefficients (Eq. 1; Fig. 1A).

### Supplementary Tables

**Supplementary Table 1. Spatial wavelengths of the first 1024 eigenmodes.**

| **Eigengroup** | **Wavelength (mm)** | **Eigenmodes included in the eigengroup** |
| --- | --- | --- |
| 0 | $-$ | 1 |
| 1 | 297.7 | 2–4 |
| 2 | 171.9 | 5–9 |
| 3 | 121.5 | 10–16 |
| 4 | 94.1 | 17–25 |
| 5 | 76.9 | 26–36 |
| 6 | 65.0 | 37–49 |
| 7 | 56.3 | 50–64 |
| 8 | 49.6 | 65–81 |
| 9 | 44.4 | 82–100 |
| 10 | 40.1 | 101–121 |
| 11 | 36.6 | 122–144 |
| 12 | 33.7 | 145–169 |
| 13 | 31.2 | 170–196 |
| 14 | 29.1 | 197–225 |
| 15 | 27.2 | 226-256 |
| 16 | 25.5 | 257-289 |
| 17 | 24.1 | 290-324 |
| 18 | 22.8 | 325-361 |
| 19 | 21.6 | 362-400 |
| 20 | 20.5 | 401-441 |
| 21 | 19.6 | 442-484 |
| 22 | 18.7 | 485-529 |
| 23 | 17.9 | 530-576 |
| 24 | 17.2 | 577-625 |
| 25 | 16.5 | 626-676 |
| 26 | 15.8 | 677-729 |
| 27 | 15.3 | 730-784 |
| 28 | 14.7 | 785-841 |
| 29 | 14.3 | 842-900 |
| 30 | 13.8 | 901-961 |
| 31 | 13.4 | 961-1024 |

**Supplementary Table 2. HCP task contrasts.**

| Task type | Number of contrasts | Contrasts | Key contrast |
| --- | --- | --- | --- |
| social | 3 | random; tom; tom_random | tom_random |
| motor | 13 | cue; lf; lh; rf; rh; t; avg; lf_avg; lh_avg; rf_avg; rh_avg; t_avg; cue_avg | cue_avg |
| gambling | 3 | punish; reward; punish_reward | punish_reward |
| working memory (wm) | 19 | 2bk_body; 2bk_face; 2bk_place; 2bk_tool; 0bk_body; 0bk_face; 0bk_place; 0bk_tool; 2bk; 0bk; body; face; place; tool; body_avg; face_avg; place_avg; tool_avg; 2bk_0bk | 2bk_0bk |
| language | 3 | math; story; math_story | math_story |
| emotion | 3 | faces; shapes; faces_shapes | faces_shapes |
| relational | 3 | match; rel; match_rel | match_rel |

**Supplementary Table 3. Specifications for testing computer.**

| **Part** | **Hardware/Specification** |
| --- | --- |
| CPU* | 2.4 GHz Quad-Core Intel Core i5 |
| L2 Cache | 256 KB (per core) |
| L3 Cache | 6 MB |
| RAM | 16 GB 2133 MHz DDR3 |
| Graphics Card | Intel Iris Plus Graphics 655 1536 MB |
| OS | macOS Catalina 10.15.7 |

* note: for all results reported in Supplementary Information-S10 and Fig-S9, we did not compute any surrogates with more than one thread per computation.
